# Flexible encoding of objects and space in single cells of the dentate gyrus

**DOI:** 10.1101/2021.06.25.449795

**Authors:** Douglas GoodSmith, Sang Hoon Kim, Vyash Puliyadi, Guo-li Ming, Hongjun Song, James J. Knierim, Kimberly M. Christian

**Affiliations:** Department of Neuroscience and Mahoney Institute for Neurosciences, Perelman School for Medicine, University of Pennsylvania, Philadelphia, PA 19104, USA; Department of Cell and Developmental Biology, Perelman School for Medicine, University of Pennsylvania, Philadelphia, PA 19104, USA; Institute for Regenerative Medicine, Perelman School for Medicine, University of Pennsylvania, Philadelphia, PA 19104, USA; Department of Psychiatry, Perelman School for Medicine, University of Pennsylvania, Philadelphia, PA 19104, USA; The Epigenetics Institute, Perelman School for Medicine, University of Pennsylvania, Philadelphia, PA 19104, USA; Department of Psychological and Brain Sciences, Johns Hopkins University, Baltimore, MD 21218, USA; Zanvyl Krieger Mind/Brain Institute, Johns Hopkins University, Baltimore, MD 21218, USA; The Solomon H Snyder Department of Neuroscience, Johns Hopkins University School of Medicine, Baltimore, MD 21205, USA; Kavli Neuroscience Discovery Institute, Johns Hopkins University School of Medicine, Baltimore, MD 21205, USA

**Keywords:** dentate gyrus, hippocampus, place cell, granule cell, mossy cell

## Abstract

The hippocampus is involved in the formation of memories that require associations among stimuli to construct representations of space and the items and events within that space. Neurons in the dentate gyrus (DG), an initial input region of the hippocampus, have robust spatial tuning, but it is unclear how nonspatial information may be integrated with spatially modulated firing at this stage. We recorded from the DG of 21 adult mice as they foraged for food in an environment that contained discrete objects. By classifying recorded DG cells into putative granule cells and mossy cells, we examined how the addition or displacement of objects affected the spatial firing of these DG cell types. We found DG cells with multiple firing fields at a fixed distance and direction from objects (landmark vector cells) as well as cells that exhibited localized changes in spatial firing when objects in the environment were manipulated. When mice were exposed to a second environment with the same objects, DG spatial maps were completely reorganized, suggesting standard global remapping, and a largely different subset of cells responded to object manipulations. Together, these data reveal the capacity of DG cells to detect small changes in the environment, while preserving a stable spatial representation of the overall context.

## Introduction

One of the hallmark features of the hippocampus in rodents is the spatially-tuned activity of its principal neurons. Place cells in the hippocampus fire whenever an animal passes through a specific location within an environment, and a place cell’s activity within its place field can be modulated by nonspatial events and information.^1–9^ An internal “cognitive map”, formed by embedding nonspatial and object-related information onto a stable spatial framework, has been theorized to promote spatial navigation in animals and episodic memory in humans.^10–13^ Although most studies on the integration of nonspatial and spatial information have focused on CA1, the dentate gyrus (DG) is the primary entry point for cortical information into the hippocampus; it receives inputs from both the medial and lateral portions of the entorhinal cortex (MEC and LEC, respectively). Distinct patterns of spatial and nonspatial information have been observed in both MEC and LEC,^14–17^ but the extent to which the DG integrates this information to support the formation of a cohesive neural representation of an experience is unclear.

In contrast to other subregions of the hippocampus, the DG contains two excitatory cell types: granule cells in the granule cell layer and mossy cells in the hilus.^18^ While most theories of DG function focus on the more numerous granule cells, mossy cells receive inputs from hundreds of granule cells and can broadly regulate granule cell activity directly and via DG interneurons.^19^ As a result, DG function relies on the coordination and communication of multiple cell types within the DG circuit.^20–23^ In behavioral studies, DG lesions do not affect the ability of an animal to discriminate among different objects, but they can impair learning of complex object-place configurations^24, 25^ or odor-context associations.^26^ Likewise, targeted optogenetic inhibition of mossy cells impairs object-place learning (with no effect on object discrimination).^21^ These results indicate that the DG may be essential for the conjunctive encoding of spatial and non-spatial signals from the medial and lateral entorhinal cortex,^27–29^ supporting the formation of complex associations.

Although these behavioral results support the conjunctive encoding of spatial and non-spatial information in the DG, it is unclear whether this information is integrated at the circuit-level among different neuronal subpopulations or within single neurons. One possibility is that DG cells are specialized for either purely spatial or object-related information. In this scenario, conjunctive representations would emerge from integrating information across the active DG population. At the other extreme of possibilities for conjunctive encoding, individual DG cells may be responsive to specific object-spatial configurations, exhibiting a unique firing pattern for each configuration of objects and cues within a particular environment such that small changes would lead to global remapping. In head-fixed mice, the addition of tactile cues to a treadmill has been shown to induce or reorganize some granule cell and mossy cell firing fields and revealed differences between the cell types in the timing and duration of cue-modulated firing during learning.^30–32^ Given the well-established ability of the hippocampus to generate highly stereotyped sequential representations of distance and time in linear tracks and treadmills, it can be difficult to disentangle neural responses to tactile cues and spatial information that is repeatedly experienced by the animal in a particular order. Therefore, it is also critical to record from freely-moving animals in a 2D environment to determine how spatial and non-spatial information is represented when not constrained by sequential presentations of stimuli and path-invariant sampling of space, which has been challenging due to technical difficulties in recording and identifying different DG cell types in vivo.

Here, we used a machine-learning approach to classify granule cells and mossy cells recorded as mice foraged for food in one or two environments with discrete objects that were manipulated in alternating sessions. Although most active DG granule cells and mossy cells maintained a stable, spatial representation of the environment with one or more place fields, many cells fired at a fixed vector relationship to objects (landmark vector cells) or significantly altered their spatial firing in response to object manipulation. Further, a complete reorganization of place field activity (global remapping) occurred between environments and object-related activity was observed in largely non-overlapping ensembles of DG cells in each environment, indicating that there is not a dedicated subpopulation of DG cells that respond specifically to objects. Together, these results indicate significant integration of distinct types of information even at the first stage of hippocampal processing and demonstrate that both spatial and non-spatial signals can be observed within the activity of single DG cells. These results also suggest flexibility within the DG population, in which many cells have the capacity to detect changes in object number and location, and can be differentially responsive to local changes in the environment, in a context-dependent manner.

## Results

### Recording from classified DG cell types

Recording sessions with two objects in standard locations (STD) were alternated with manipulation (MAN) sessions in which one object was moved (MOVE) or a third object was added (ADD) (Figure 1A-C). Of 366 well-isolated cells recorded from the DG hilus and granule cell layer (Figure 1D), 178 (49%) did not have a significant place field in any recording session. The remaining 188 cells (51%, “active cells”) had at least one place field in at least one session. There were between 2 and 8 recording sessions for each cell, with a total of 1513 recording sessions across the 366 cells: 923 STD sessions and 590 MAN sessions (160 ADD and 430 MOVE sessions). There was no difference between the proportions of STD and MAN sessions that had at least one place field in the environment (sessions with ≥ 1 field: STD 415/923, MAN 261/590; χ(1)^2^ = 0.08, p = 0.78), indicating that manipulation of objects within the environment did not cause a significant change in the proportion of cells that were active. Most of the analysis presented here will focus on the 188 active cells with at least one place field in at least one recording session.

**Figure 1:**
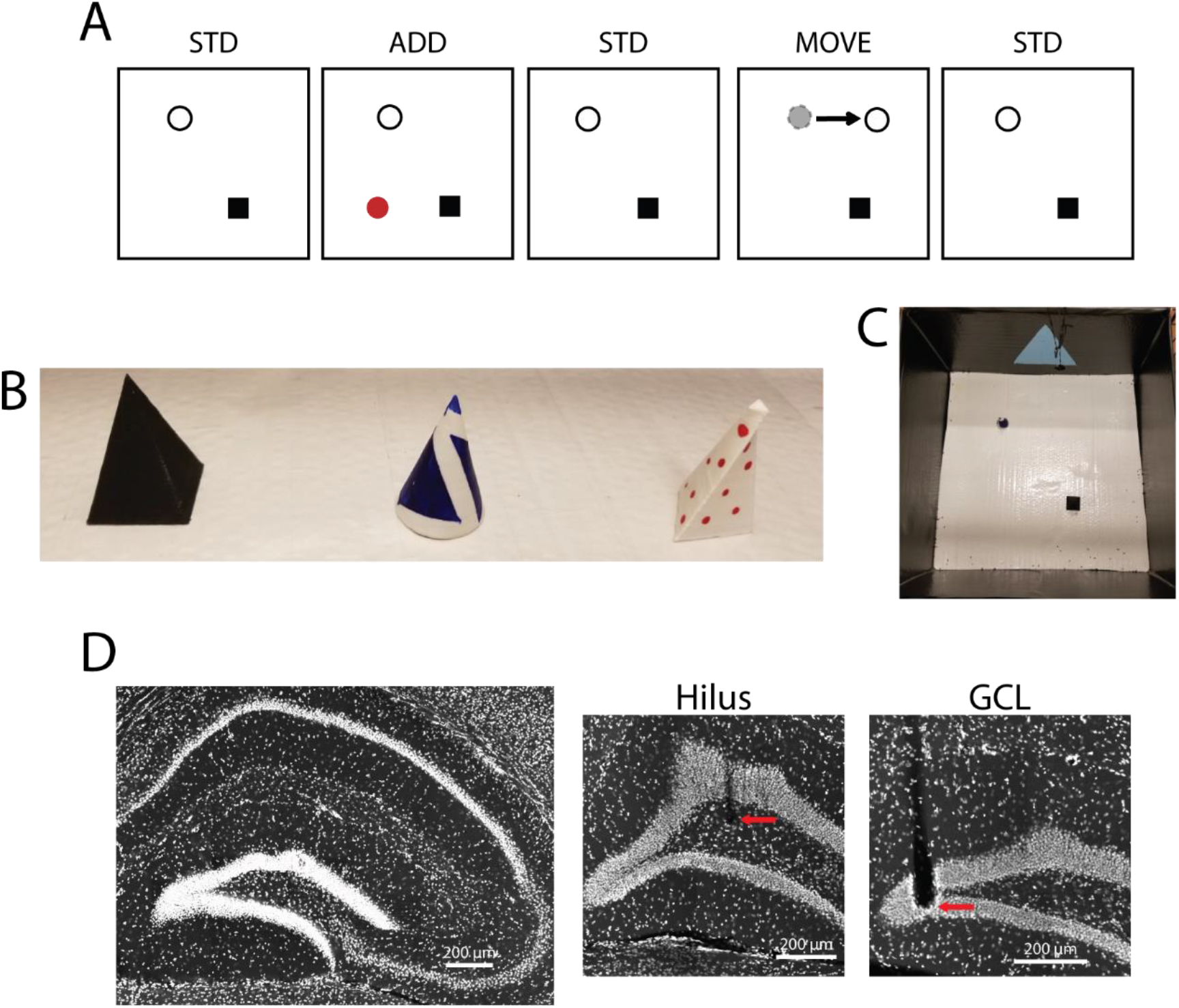
Behavioral task and DG recordings. A) Schematic of example recording day. Standard (STD) sessions in which two objects were in a standard location were alternated with manipulation (MAN) sessions, in which a third object was added (ADD) or one object was moved (MOVE). B) Picture of objects used, the two objects on the left are the STD objects. C) Picture of the environment with objects in the STD configuration. D) Image of the hippocampus (left) and higher magnification images of tetrode tracks terminating in the hilus and granule cell layer (GCL).

Mossy cells can be recorded on electrodes located up to 300 µm away and are much more active than granule cells.^33–36^ The majority of active cells recorded extracellularly in the DG are, therefore, likely to be mossy cells.^22^ For this reason, it is more difficult to infer cell identity based on the recording site and alternative methods should be used to determine cell-type specific firing properties of granule cells and mossy cells. In previous work, we showed how machine learning techniques could be used to resolve the activity of these distinct DG cell types.^22, 33^ To address how object-related firing changes may be distributed across granule cell and mossy cell populations, we utilized similar methods to assign putative cell types to cells recorded in the DG of mice. We trained a random forests classifier to separate these cell types based on their firing properties in a post-behavior rest/sleep session (see STAR Methods, Figure 2). The training data for this classifier was a group of DG cells recorded in a separate foraging task (recorded in the same mice but on different recording days; see STAR Methods, Figure 2A). Based on prior studies of DG cell activity,^33, 34^ granule cells and mossy cells were identified in the training data based on the number of distinct environments in which the cell had place fields, and the classifier was trained to separate between these cell types based on firing features recorded in a post-behavior sleep session (Figure 2A). The classification features were the mean firing rate, burst index, spike duration, and the first principal component of the second derivative of the waveform^33, 34, 36^ (Figure 2B). The classifier reliably discriminated between cell types and had an estimated error rate of ∼5% (see STAR Methods, Figure S1). Importantly, the classifier identified DG cell types on object recording days using firing properties recorded in the post-behavior sleep session only (no behavior sessions). Of the 366 cells recorded in the DG, 172 cells were classified as mossy cells and 194 cells were classified as granule cells. 83% of cells without fields (147/178) were classified as granule cells, and 75% of active cells (141/188) were classified as mossy cells. Compared to classified mossy cells, classified granule cells (active cells) had fewer place fields (granule cells median 1, IQR 1 – 2; mossy cells median 2, IQR 1 – 2; rank-sum test z = 3.75, p = 1.78 x 10^-4^), lower mean firing rates (granule cells 0.70 ± 0.04 Hz; mossy cells 1.03 ± 0.03 Hz; rank-sum test z = 5.52, p = 3.31 x 10^-8^), and lower peak firing rates (granule cells 6.73 ± 0.52 Hz; mossy cells 11.98 ± 0.36 Hz; rank-sum test z =3.98, p = 6.93 x 10^-5^) in behavior sessions. These results are consistent with previous reports of the firing properties of DG cell types^33, 34, 37, 38^, further supporting the reliability of our classification method and demonstrating the utility of these classification methods for analyzing DG extracellular recordings.

**Figure 2:**
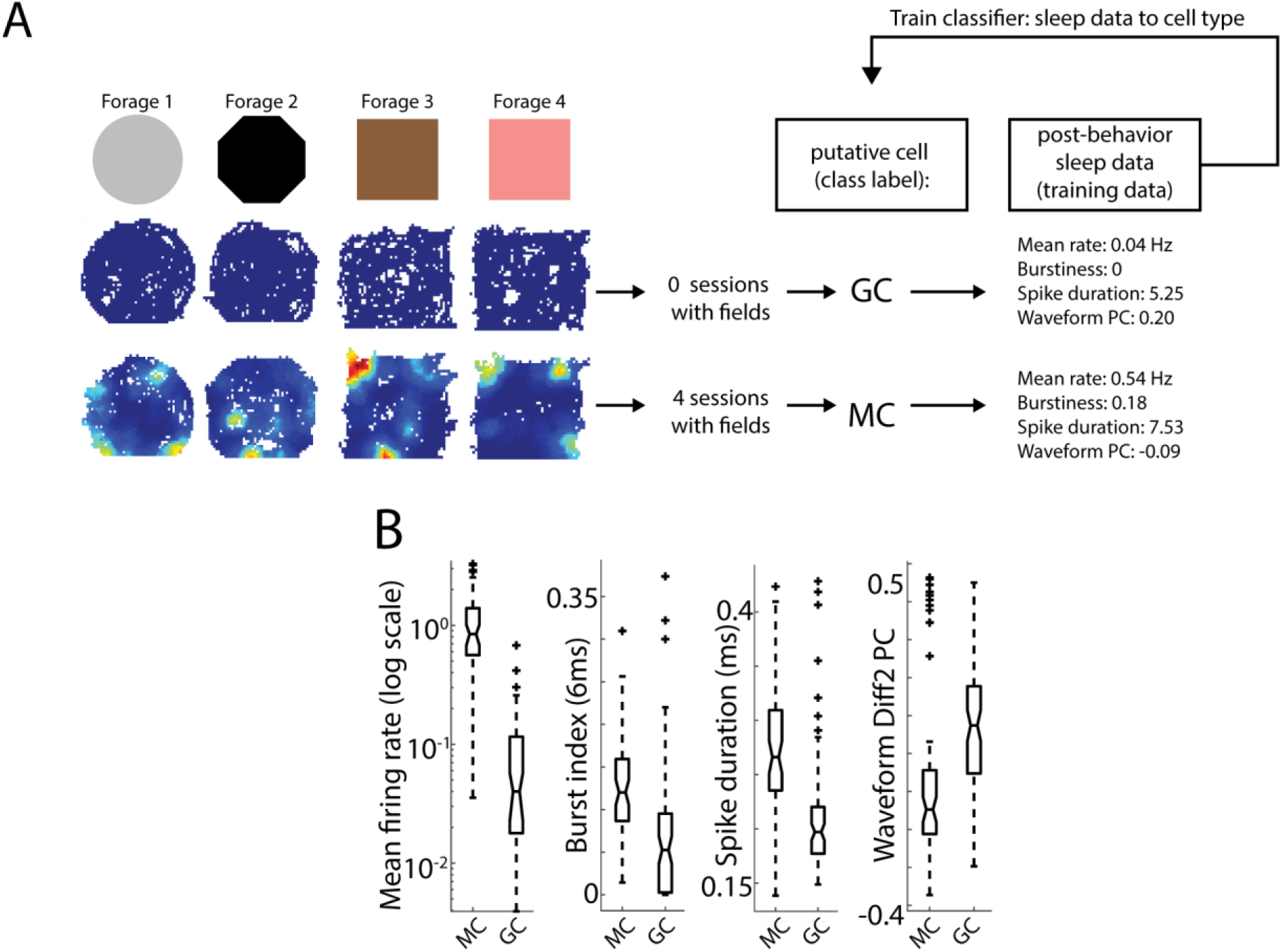
Classification of DG cell types. A) Cells recorded in a foraging task (without objects) were used to train a classifier to separate putative granule cells and mossy cells. DG cells were recorded as animals foraged for food in four distinct environments. Based on previous reports of DG place fields^33, 34, 37, 68^ most granule cells should have no place fields in any environment, and most mossy cells should have place fields in most environments. We therefore considered any cell with no fields in any session as a putative granule cell and any cell with fields in 3 or more sessions as a putative mossy cell. After these putative cell type labels were assigned, the properties of these cells in a post-behavior sleep session were identified. These firing features were used as the training data, and a random forests classifier was trained to classify the training data to the putative cell type labels. B) Features used for classification. As classification features are correlated with cell type, these plots are presented for display purposes only and statistical comparisons were not performed. Boxplots indicate the values for putative granule cells (GC) and mossy cells (MC) in our training data. The dashed line indicates the range of data (outliers marked +), horizontal lines indicate the median, 25th percentile, and 75th percentile, and notches indicate the 95% CI of the median. From left to right: mean firing rate, burst index, spike duration, and first principal component of the second derivative of the waveform.

### Population activity and place field comparisons

We first aimed to determine the effect of object manipulation on overall DG activity levels by comparing the firing rates and place field properties of active DG cells in STD and MAN sessions. There was no significant difference in the median values of the mean firing rate between the STD and MAN sessions for active granule cells or mossy cells (Table S1 for statistics; Figure 3A). Similarly, the median values of peak firing rates in STD and MAN sessions were not significantly different (Table S1 for statistics; Figure 3B).

**Figure 3:**
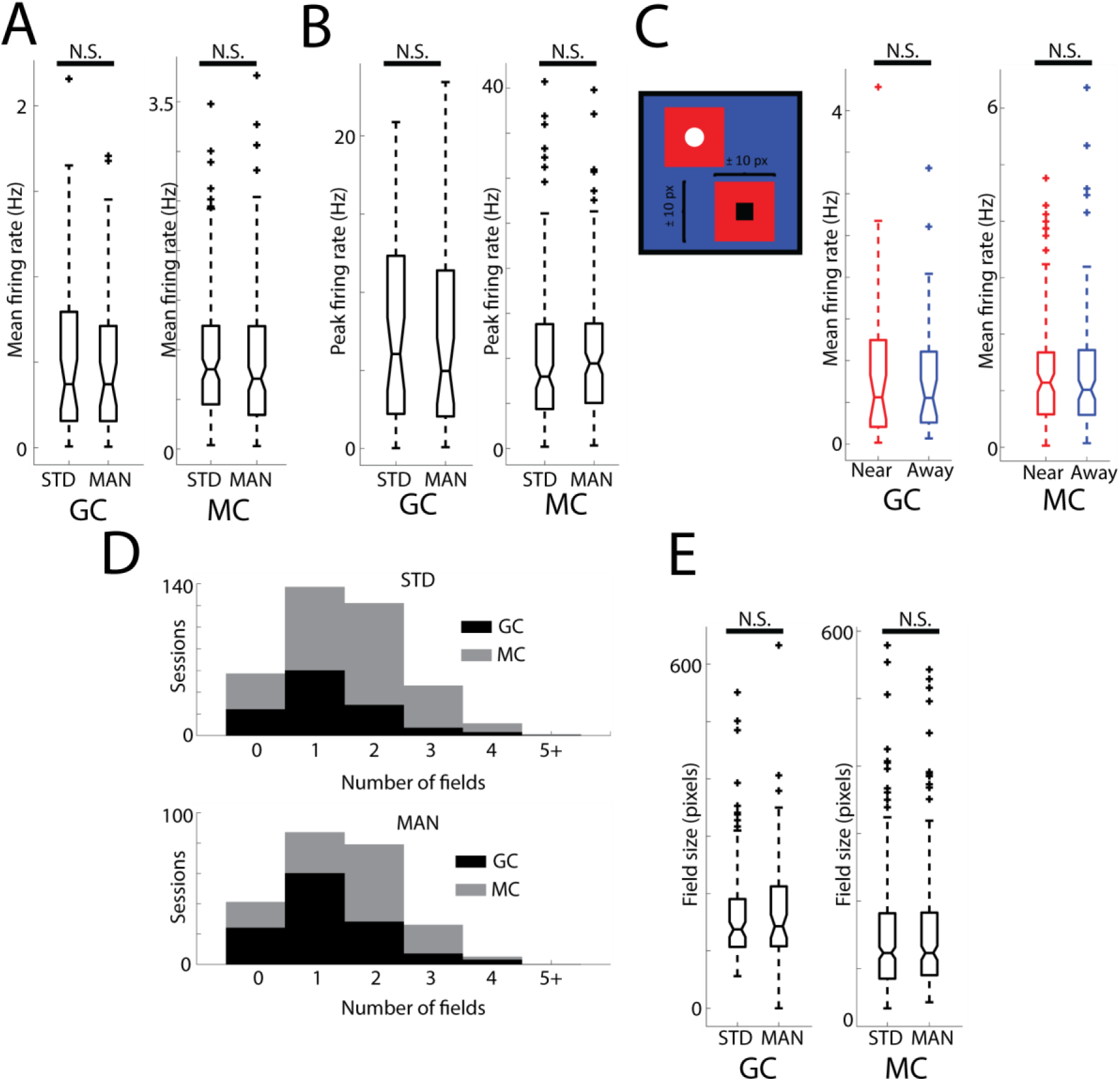
DG population firing rate and place field properties in STD and MAN sessions. A) Median value of mean firing rates in STD vs. MAN sessions for granule cells (left, GC) and mossy cells (right, MC). Results of signed-rank tests are marked above, N.S. = not significant, p > .05. B) Same as in A but for peak firing rate. C) Left: mean firing rate near (red area, ± 10 pixels from an object location) or away from (blue area, > 10 pixels from all objects) objects. Right, same as in A for mean firing rate near vs. away from objects for all sessions. D) Number of place fields for active cells (place field in at least one session). There was no significant difference in the number of fields in STD (top) vs. MAN (bottom) sessions for either granule cells (black) or mossy cells (gray). Granule cell histograms are plotted to overlay mossy cell histograms. E) Same as in A but for place field size (in pixels).

Although there were no differences in overall firing rates following the manipulation of objects in the environment, changes in firing rates may have occurred only when the animal was close to an object. To determine if DG cells were more likely to fire close to an object than farther away from an object, we compared the average firing rate within 10 pixels of an object to all other pixels. There was no significant difference between the firing rates of DG cells near objects and their firing rate > 10 pixels away from all objects (Table S1 for statistics; Figure 3C).

If the manipulation of objects generally caused the formation of new place fields or the elongation of place fields, more numerous or larger place fields would be expected in MAN sessions than STD sessions. However, there were no differences in the number of place fields in STD vs. MAN sessions for cells active in at least one session (Table S1 for statistics; Figure 3D) or in the average place field size (Table S1 for statistics; Figure 3E). It is possible that the addition of objects, specifically, could cause changes in the firing rates or place field properties of DG cells. However, no significant differences in these properties were observed when restricting our analysis to ADD sessions (Table S1). Together, these results demonstrate that the addition and manipulation of objects within the environment did not significantly and coherently alter firing rates or place field size/number across the granule cell or mossy cell population.

### Landmark vector cells

Although we did not detect any significant population differences in firing rate or place field size/number, visual inspection of the rate maps revealed many DG cells that appeared to have firing related to the presence of objects within the environment. Some of these observed responses were consistent with that of previously described landmark vector (LV) cells. LV cells have been recorded in the hippocampus^4, 5^ and MEC^17^ (referred to as object-vector cells in MEC^17^). These cells fire at a fixed vector relationship (same distance and direction) to objects/landmarks within the environment, and many DG cells in the present study appear to have this property. In order to identify LV responses in an individual recording session, it is necessary to analyze cells with at least two firing fields (i.e., a cell with a single field near an object could be a LVC, but it could also be a cell with a spatial place field that happens to be near an object). Putative LVCs were identified based on the minimum pairwise difference between vectors connecting objects to place field centers^4^ (see STAR Methods, Figure 4A).

**Figure 4:**
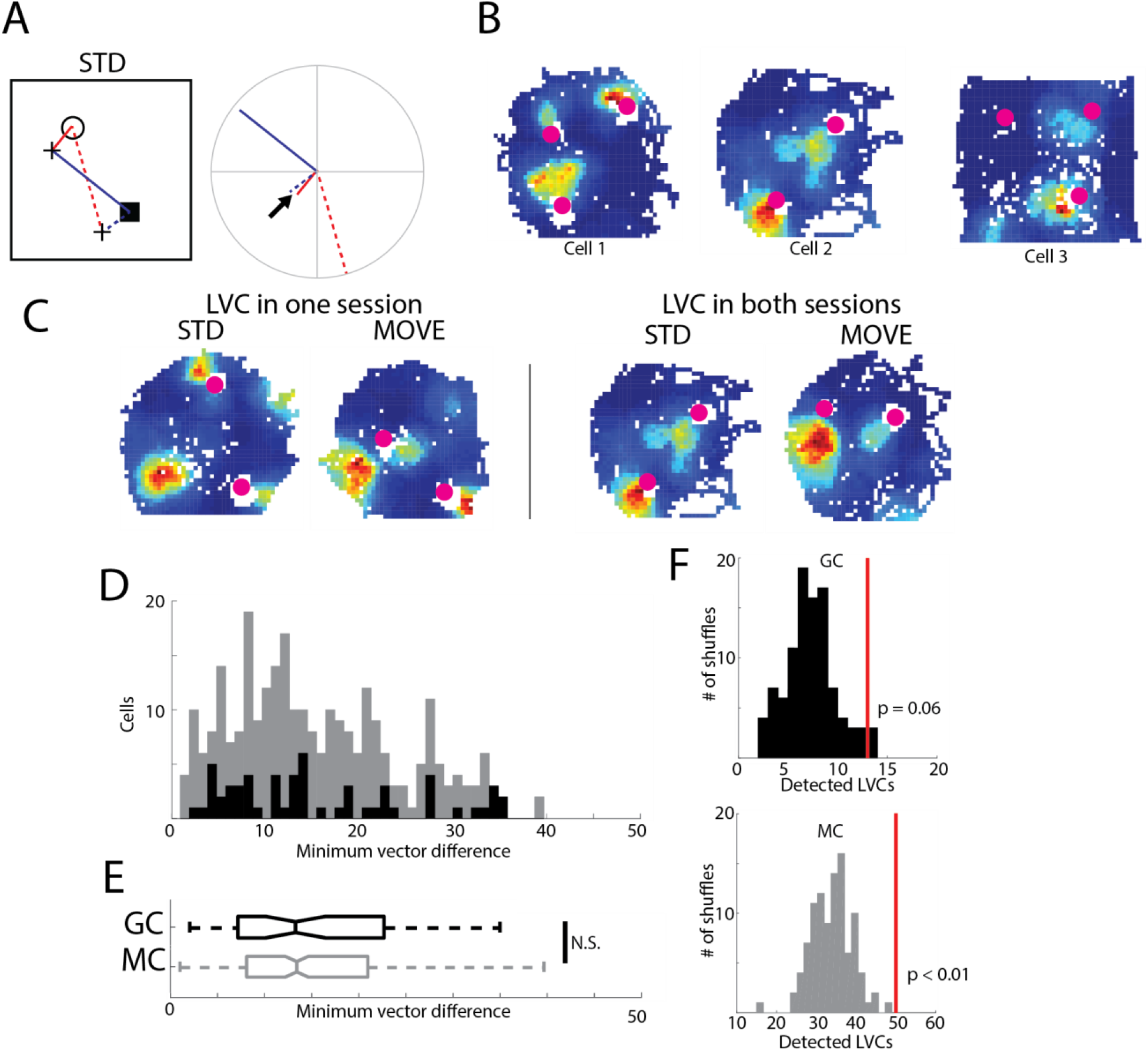
Landmark vector responses. A) Schematic of the procedure for identifying putative LV responses in a single session. On the left, the vectors between each object (circle and square) and each place field centroid (black “+”) are determined (red/blue solid/dashed lines). On the right, the distance and direction from the object is plotted for all vectors (center is object location). The difference is calculated between all pairs of vectors that do not share the same object (same line color) or same place field centroid (same line style). The pair of vectors (that do not share an object or place field) with the smallest difference between them is determined (black arrow). If this minimum vector difference is < 7 pixels, the response is considered a LV response. B) Example sessions in which the minimum vector difference was < 7 pixels (putative LV responses). C) Example session pairs (STD and MOVE) detected as LV responses in only the MOVE (left) or both (right) sessions. D) Distribution of observed minimum vector difference values for granule cells (black) overlaid over values for mossy cells (gray). E) Boxplot of data from D. F) Random distributions of minimum vector difference values were generated 100 times by randomly assigning place field centers to each session; the number of detected LV responses in each of these 100 random distributions is plotted for granule cells (top, black) and mossy cells (bottom, gray). The red line indicates the number of LV responses observed in our data.

LV responses were detected in 37 mossy cells and 10 granule cells (see Figure 4B-C for examples). LV responses in some cells were only observed in a single session while LV responses were present across multiple sessions in other cells (Figure 4C). LV responses were observed in more than one session for 8/37 mossy cells and 3/10 granule cells. However, mossy cells were more likely than granule cells to have the multiple place fields required to detect a LV response in a session. Of the 331 sessions with at least two place fields, putative LV responses were identified in 18% (50/273; significantly greater than the 5% expected by chance; test for proportions z = 10.10, p = 2.93x10^-24^) of classified mossy cell sessions and 22% (13/58; test for proportions z = 6.09, p = 5.82x10^-10^) of classified granule cell sessions. There was no significant difference between cell types in the proportion of LV responses detected in sessions with at least two fields (χ(1)^2^ = 0.52, p = 0.47). There was also no significant difference in the distribution of minimum vector differences between granule cells and mossy cells (granule cells median 13.49, IQR 7.28 – 23.01; mossy cells median 13.43, IQR 8.11– 20.97; rank-sum test z = 0.21, p = 0.83; Figure 4D, E).

Observed minimum vector differences were then compared to values obtained by randomizing place field locations (see STAR Methods, ref ^4^). The procedure for generating random granule cell and mossy cell distributions was repeated 100 times, and the number of observed LVCs was larger than all random distributions for mossy cells (see STAR Methods, p < 0.01), but granule cells only showed a statistical trend in that direction (p = 0.06) (Figure 4F), even when a stricter threshold was used for LV detection (Figure S2). These results indicate that there were more LV responses than expected by chance in mossy cells, with less convincing evidence for granule cells. In summary, a significant number of LV responses were detected in the DG. LV responses were detected in more sessions than expected by chance in mossy cells, while granule cells showed only a trend in that direction. However, the proportion of sessions with a detected LV response did not significantly differ between the cell types.

### Diverse DG cell responses to object manipulation

In addition to the LV responses, which are consistent with previous reports in the hippocampus and entorhinal cortex and can be observed in a single session,^4, 5, 17^ manipulation of objects across sessions allowed us to detect a considerable array of additional object-related changes in DG cell activity (Figure 5A). Overall, apparent object-related firing changes could be divided into at least six different response categories (Figure 5B-G). “Moved field” responses (cells with firing fields located near an object both before and after that object was moved) frequently appear in our recordings (moved field; Figure 5B). These fields would often maintain their vector relationship with the moved object, similar to LV responses. Other times, however, a field would maintain its distance to the moved object but rotate to a different angle relative to the object (rotation; Figure 5C). These responses likely did not represent a global rotation of the rate map, as individual fields often rotated around an object while the firing locations of other fields of the same cell were unchanged (Figure 5C).

**Figure 5:**
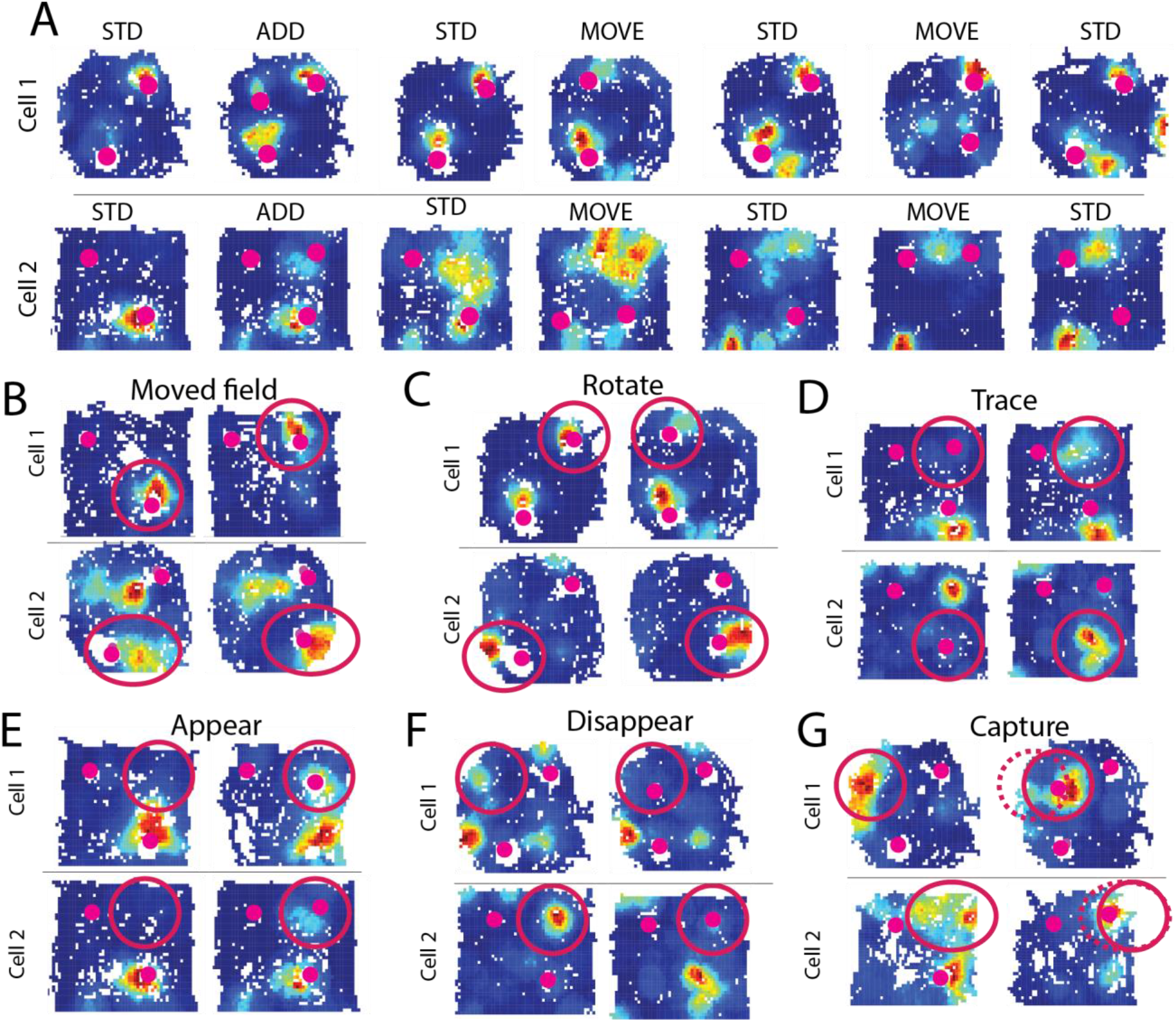
Example object-related DG activity changes. A) Two example cells with seven recording sessions. Each row is one cell, and the rate map for seven consecutive sessions is shown. Object locations are represented by magenta dots. Diverse object-related changes in activity can be identified in these cells. Cell 1 has multiple fields with the same vector relationship to objects in the first three sessions (putative LV responses). In the next session, a field moves with the moved object but rotates to fire at a new angle relative to the object. In the same session, a new field begins to form near the unmoved object. When the second object is moved, there are weak fields at both the current and previous location of that object. In cell 2, a new field forms at the location of an added object, and firing persists at that location throughout the remaining sessions. The stable firing field near the bottom object disappears when that object is moved. B-G) Observed types of object-related activity changes. Two example cells are shown for each type of response, and two consecutive sessions are shown for each cell. Red circles indicate the field that demonstrates the object-related response. B) Moved field response. C) Rotate response. D) Trace response. E) Field appears. F) Field disappears. G) Capture (the dashed red line indicates the prior location of the field).

We also observed “trace cells,” which do not fire strongly at the location of an object while the object is present but fire at the location where a moved object had previously been^4, 39, 40^ (trace; Figure 5D). These responses do not reflect the current relationship between objects and the environment, and instead reflect some “memory” of a previous object configuration. Other cells had place fields appear at the location of a new or moved object (appear; Figure 5E) or disappear when an object was placed near the field location (disappear; Figure 5F). Finally, “capture” responses occurred when an existing place field moved closer to a new or moved object (capture; Figure 5G).

Object-related firing was only observed in a subset of DG cells; most cells and individual place fields displayed stable spatial firing patterns (Figure S3), as expected from hippocampal place fields. Intriguingly, individual place fields of the same cell were often differentially affected by object manipulations; some fields would appear to respond to object manipulations while other fields of the same cell remained stable at the same spatial location (Figure S3). This pattern of activity indicates that DG cells are not specialized for either spatial or object-related firing, but rather suggests that individual fields of a DG cell can exhibit varied responses to objects ranging from a stable spatial field with no object-related modulation (Figure S3) to a highly modulated field responsive to object manipulations (Figure 5, S3). In addition, different types of object responses, including LV responses, could often be observed in the same cell across sessions (Figure 5A, S3). The presence of multiple object-related activity changes within the same cell indicates that object response types are not fixed; they represent a dynamic and flexible response to object manipulations within the DG population.

### Identification and characterization of putative object-responsive DG cells

Given that individual fields could be differentially responsive to object manipulation, we next asked whether we could identify sessions in which the location of place fields changed significantly in the MAN sessions. Many of the response types observed reflected a place field that was closer to the final location of the moved object (X in Figure 6A, top) in the MAN session than in the STD session (i.e., MOVE, ROTATE, CAPTURE, APPEAR responses). Therefore, we first identified sessions in which the distance between the location of the manipulated object (either moved or added) and the nearest field was significantly reduced in MAN sessions (Figure 6A, see STAR Methods). While multiple session pairs could come from the same cell, different session pairs reflect distinct object manipulations (no object manipulations were repeated on the same day). The distance from the manipulated object to the nearest place field decreased more than expected by chance (see STAR Methods, Figure 6A) for a significant number of mossy cell session pairs (17 out of 157 pairs, 11%; test for proportions z = 3.35, p = 4.03x10^-4^) but not for granule cell session pairs (4 out of 54 session pairs, 7%; test for proportions z = 0.81, p = 0.21) (Figure 6A). However, there was no significant difference in the proportion of detected session pairs between cell types (χ(1)^2^ = 0.52, p = 0.47) The majority of detected session pairs came from unique cells (15 unique mossy cells and 4 unique granule cells). These results indicate that manipulating objects causes a change in the location of firing fields within a significant subpopulation of DG cells.

**Figure 6:**
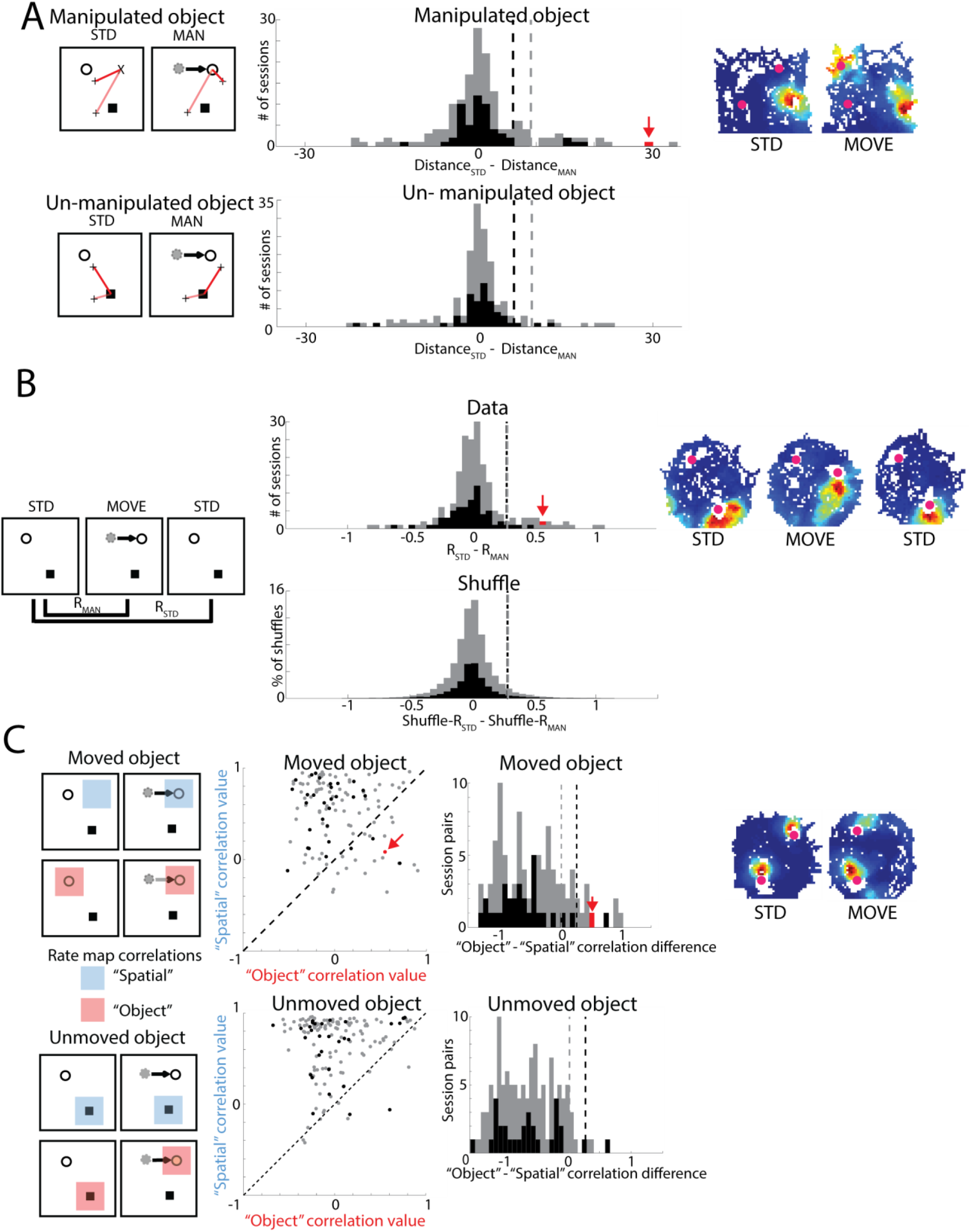
Detection of cells with responses to object-manipulation. A) Distance to nearest place field centroid. Left: Schematic of distance calculation in a pair of STD and MAN sessions. The distance (length of the red line) from the location of the moved object (in the MAN session, “X” in STD, black circle in MAN) to each place field centroid (black “+”) was determined. The shortest distance in each session was determined (dark red line), and the difference between these distances was calculated. Middle: The distance between the manipulated object location and all place fields (top) was calculated and compared to the distribution of distances between the un-manipulated object and all place fields (bottom). The dashed lines reflect the 95^th^ percentile of the un-manipulated object distributions. Right: spatial firing of an example session pair (marked in red on the histogram) with a significant reduction in the distance between object location and place field centroid. B) Rate map correlation difference. Left: Schematic of three sequential sessions (STD, MAN, and STD). The correlation between STD sessions was calculated (R_STD_), as well as the correlation between the first STD and the MAN session (R_MAN_). Middle: Distribution of correlation difference values between R_STD_ and R_MAN_. On the bottom is the distribution generated by randomly shuffling the identity of the first STD session 100 times and calculating correlation difference values. Cells with a higher correlation difference than the 95th percentile of the shuffled distributions (dashed lines) were considered significant. Right: spatial firing of an example of three consecutive sessions with a significant correlation difference value (marked in red on the histogram). C) Local rate map correlation difference. Top: moved object, bottom: un-moved object. From left to right: schematics of the location of “object” and “spatial” rate map correlations, scatter plot of “object” vs. “spatial” correlation values, distribution of the difference between “object” and “spatial” correlation values, example of session pair with detected object response (marked in red on histogram and scatter plot). For all plots, mossy cell data is presented in gray and granule cell data is presented in black.

While the preceding analysis can detect multiple types of responses to object manipulation, it is not sensitive to responses that do not result in firing proximal to objects (i.e., DISAPPEAR, TRACE). Therefore, we next examined whether manipulation of objects caused detectable changes in spatial rate map correlations. In any sequence of STD – MAN – STD sessions (Figure 6B), the rate map correlation between STD1 and STD2 (R_STD_) was expected to be high, as the environment and object locations were identical in these sessions. Changes in the rate map in the MAN session, however, were expected to reduce the correlation between the STD1 and MAN session rate maps (R_MAN_). Across the active DG population, there was no significant difference between the median R_STD_ and the median R_MAN_ (R_STD_ 0.67 ± 0.02; R_MAN_ 0.65 ± 0.02; signed-rank test z = 1.90, p = 0.06). However, the distribution of differences between R_STD_ and R_MAN_ correlations reveals a subset of cells with a much higher STD1/STD2 correlation than STD1/MAN correlation (Figure 6B), indicating cells in which object manipulation affected spatial firing. The difference between R_STD_ and R_MAN_ exceeded the 95^th^ percentile of a shuffled distribution (see STAR Methods, Figure 6B) for a significant number of classified mossy cells (23 out of 190 session groups, 12%, test for proportions z = 4.49, p = 3.50 x 10^-6^), but not for granule cells (4 out of 63 session groups, 6%, test for proportions z = 0.49, p = 0.31; Figure 6B). There was, however, no significant difference in the proportion of mossy cell and granule cell session groups that exceeded the 95^th^ percentile of their respective shuffled distributions (χ(1)^2^ = 1.64, p = 0.20), likely due to the small number of active granule cells used for this analysis. All session groups that exceeded the 95^th^ percentile of the shuffled distribution came from unique cells. Together, these results demonstrate that more DG cells than expected by chance have reduced correlations following the displacement or addition of an object, reflecting object-related DG cell activity.

Based on the initial inspection of the rate maps, many cells have individual fields that appear to respond to object manipulations, as well as other place fields that appear unaffected by the object manipulation. These stable place fields could produce high rate-map correlation values, potentially obscuring local object-related activity when considering the overall spatial correlation between sessions. To address this, we next focused our analysis on the local area around each object. By comparing 21 x 21 pixel segments of firing rate maps based on a fixed spatial area or centered around a manipulated object, we calculated “spatial” and “object-related” correlation values, respectively (see STAR Methods). A stable and purely spatial place cell would generate a high “spatial” correlation value but a low “object-related” value. Alternatively, a cell with a place field that moved with an object would have a significantly higher “object-related” than “spatial” correlation value. There was a much higher “spatial” than “object-related” correlation value for the majority of cells, indicating the presence of stable spatial place fields. A subset of cells, however, had higher “object-related” than “spatial” correlation values (24/124 session pairs; Figure 6C, middle), suggesting spatial activity was altered by object manipulation. The majority of session pairs with a higher “object-related” than “spatial” correlation value belonged to classified mossy cells (21/24, Figure 6C, middle). However, the difference between “object-related” and “spatial” correlation values was not significantly higher for mossy cells than granule cells (granule cell median -0.63, IQR -0.93 – -0.40; mossy cell median -0.53, IQR -0.94 – -0.08; rank-sum test z = 0.1.40, p = 0.16; Figure 6C, right).

To determine how much of the difference in correlation values could result from random remapping or drift of place fields, we next calculated local rate map correlations centered around the “unmoved” object (Figure 6C, see STAR Methods). The correlation difference values were significantly higher (more “object-related”) for the “moved” object than the “unmoved” object for mossy cells (moved median -0.53, IQR -0.94 - -0.08; unmoved median -0.668, IQR -1.0 - - 0.27; rank-sum test z = 2.57, p = 0.01) but not granule cells (moved median -0.63, IQR -0.93 – - 0.41; unmoved median -0.67, IQR -1.05 – -0.19; rank-sum test z = 0.23, p = 0.82). This result suggests that the population activity of mossy cells, but not granule cells, was significantly altered by object manipulations. However, as the correlation values for the moved object did not significantly differ between granule cells and mossy cells, this difference likely reflects the effect of sparse granule cell firing on the “unmoved” control correlation values. Of the 30 active session pairs from classified granule cells, 1 (3%) had a correlation difference value that exceeded the 95^th^ percentile of the “unmoved” object distribution (test for proportions z = -0.42, p = 0.66). For classified mossy cells, however, 19 (from 18 unique cells) of the 94 active session pairs (20%) had a correlation difference value higher than the 95^th^ percentile of the “unmoved” distribution (test for proportions z = 6.77, p = 6.55 x 10^-12^; Figure 6C, right), indicating that object manipulation altered the activity of significantly more mossy cells than expected by chance. The proportion of cells significantly affected by object manipulation (correlation difference exceeded the 95^th^ percentile of “unmoved” distribution) was significantly higher for mossy cells than granule cells (χ(1)^2^ = 4.79, p = 0.03).

### DG object responses in distinct environments

Following object manipulations, the overall DG spatial representation was largely unchanged, and firing fields of a subset of DG cells were modified. We next aimed to determine how exposure to distinct environments would affect the object-related activity of DG cells. To determine whether cells with object-related activity in one environment would respond similarly in a different environment, we recorded from 3 additional mice as they foraged for food in two environments containing objects (ENV_A_ and ENV_B_, see STAR methods, Figure 7). The two environments had the same size and shape but different walls and floors, and the objects, object locations, and object manipulations were identical in each environment (Figure 7A).

**Figure 7:**
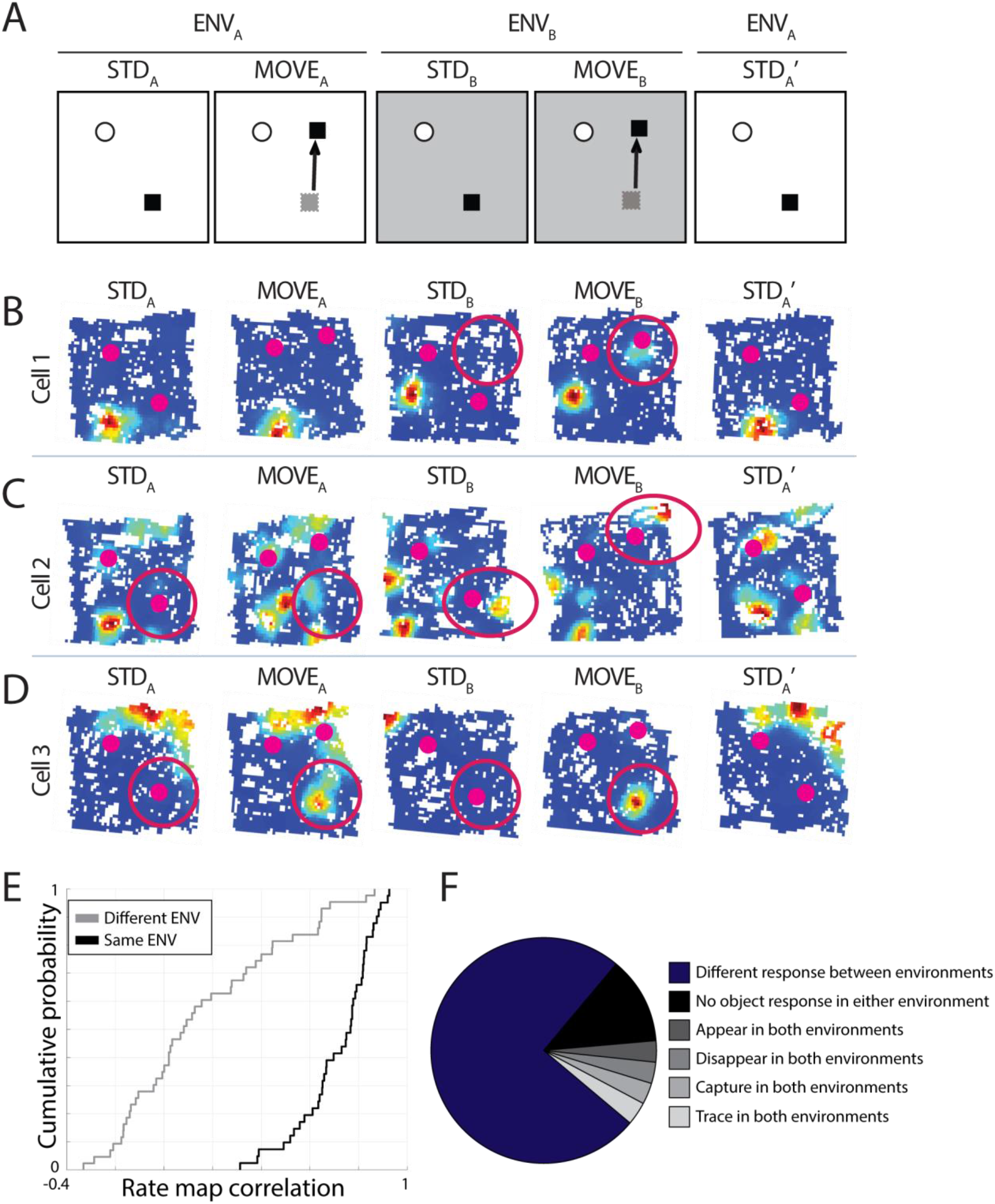
Recordings from two distinct environments. A) Schematic of recording day. A STD and MOVE session in one environment (ENV_A_) were followed by a STD and MOVE session in a second environment (ENV_B_). The objects, their locations within the environment, and the manipulation were the same in both environments. The day ended with a final STD session in ENV_A_. B) Example cell that had no object response in ENV_A_ but had a field appear in the MOVE session of ENV_B_. C) Example of a cell with different object responses in both environments. This cell had a trace field in ENV_A,_ and a moved field in ENV_B_. D) Example of a cell with the same object-related activity in both environments. This cell had a trace response in both ENV_A_ and ENV_B_. E) Correlation between STD sessions in the same environment (STD_A_ vs. STD_A_’) or different environments (STD_A_ vs. STD_B_). Despite both environments having the same size, the same objects, and the same object locations, the correlation between different environments was significantly lower than the correlation between repeated STD sessions in the same environment, suggesting global remapping of DG activity between environments. F) Proportion of all 32 cells that were active and had unambiguous activity in both environments. The assigned response to object manipulation was the same in both environments for 8 cells. 4 cells did not change their activity following object manipulation (spatial firing appeared stable) and four had the same object-related change in activity detected in both environments (Figure S4B). The remaining cells had different responses to object manipulation in the two environments.

As the number of active granule cells is low and separate analysis of granule cells and mossy cells did not yield significantly different results, we have presented the analysis of all DG cells (combined granule cells and mossy cell data) here. We recorded from 57 DG cells that were active in at least one session through the day. There were 95 pairs of STD and MOVE sessions in which at least one field was detected (50 from ENVA and 45 from ENVB). The correlation between STD sessions across the two environments was significantly lower than the correlation between repeated STD sessions in the same environment (STD_A_ v. STD_B_ 0.16 ± 0.05; STD_A_ v. STD_A_’ 0.71 ± 0.02; rank sum test z = 6.61, p = 3.76 x 10^-11^; Figure 7E), even after rotating one rate map in 90° increments (to check for rate map rotations between environments, Figure S4A). These results indicate robust global remapping of DG cells’ spatial firing between these contexts. While global remapping of the DG population was observed between environments, object manipulation within a single environment led to local perturbations of spatial firing patterns as expected from the observations in single environments above. Although individual DG place fields were affected by object manipulations (Figure 7C, D), global remapping of all fields was only observed in response to changes in the overall environment, reflecting a stable spatial map of each environment at the population level. We next examined whether the same cells exhibited object-related DG activity in both environments to determine if there was a fixed population of cells with the capacity to respond to objects.

To evaluate object-related activity changes within each environment, three observers (D.G., S.H.K, and K.M.C) made a subjective assessment of the response to object manipulation (see STAR Methods). At least two observers identified the same object-related response type in 92% of the 95 sessions (all three selected the same response for 53% of sessions), indicating reliable identification of object responses. Of all 87 active, unambiguous (at least two observers identified the same response) sessions, 43% had spatial firing but no clear object response and the remaining 57% of sessions were split relatively evenly among other response types (7% move/rotate, 16% appear, 16% disappear, 8% trace, 10% capture). Of 32 cells with at least one firing field in both environments (excluding cells with ambiguous responses), object-related activity changes could be observed in only one of the two environments (Figure 7B) for 18 cells (56%). In 10 cells (31%), activity changes occurred in both environments after the object was moved. Among the latter group, either a different (6 cells, Figure 7C) or the same (4 cells, Figure 7D, Figure S4B) type of object response was observed in the two environments. An additional 4 cells (12%) were active in both environments but had no object-related activity change in either environment (Figure 7F). The proportion of cells with an object response in both environments did not significantly differ from the proportion expected by chance, nor did the proportion of cells with the same type of object response in both environments (Table S2). Together, these results indicate that the likelihood of an individual DG cell showing changes in object-related activity following object manipulation is independent in distinct environments, suggesting that dedicated DG populations of “object cells” and “spatial cells” are not necessary to convey information about nonspatial cues within a spatial context. Instead, these results suggest that object-related DG activity can be context-dependent and that different subsets of cells can be object-responsive in any given environment.

## Discussion

We recorded from DG granule cells and mossy cells as mice freely explored 2D environments to determine the extent to which spatial firing properties could be altered by the presence and manipulation of objects. While most cells maintained a stable, spatial map of the environment, the activity of many DG place cells was affected by the addition or displacement of objects within the environment. Many cells resembled LV cells previously reported in the hippocampus (and similar object-vector cells in entorhinal cortex),^4, 17, 30^ with multiple firing fields that shared the same vector relationship to multiple objects. Other cells had place fields that moved when a nearby object was moved, fields that appeared, disappeared, rotated, or moved closer to an object, as well as trace firing at previous object locations. When animals were placed in a second environment, global spatial remapping occurred across the population of DG cells, and cells that responded to object manipulations in one environment were often unaffected by the same manipulation in the other environment. These results suggest that many DG cells harbor the capacity to flexibly respond to object manipulations while the DG population can maintain stable spatial representations of distinct environments.

### Population-specific responses in the DG

Granule cells receive inputs from both the LEC and MEC, which are often viewed as providing non-spatial and spatial information (respectively) to the hippocampus.^40, 41^ Thus, object-related granule cell activity may be primarily driven by LEC inputs. However, studies have identified egocentric^15^ and spatial representations in LEC^39, 40^ and object vector cells in MEC^17^, suggesting each region could provide both spatial and non-spatial information to granule cells. Selective inhibition of MEC or LEC inputs to the DG could reveal the specific MEC and LEC contributions to object-related granule cell activity. In contrast, mossy cells do not receive extensive entorhinal inputs.^42^ The source of object-related activity in mossy cells is, therefore, unclear but presumably is inherited from granule cells or through CA3 backprojections.^43^ Immature adult-born granule cells (abGCs) receive preferential input from LEC,^44, 45^ and are more excitable and active than the more numerous mature granule cells (mGCs).^46–48^ These excitable abGCs may amplify object-related signals from LEC and relay this information to mossy cells. Intriguingly, abGCs have also been found to monosynaptically inhibit mGCs in response to stimulation of LEC inputs and to activate these cells following MEC input.^49^ Therefore, object-related inputs from LEC to abGCs may have the simultaneous effects of passing an amplified nonspatial signal to mossy cells while inhibiting output from mGCs.

A large majority of active cells recorded extracellularly in the DG are likely to be mossy cells.^22, 36^ Most of the active cells recorded in the present study (75%) were classified as mossy cells, and mossy cells often had multiple place fields within the environment. We found a significant number of mossy cells, but not granule cells, that responded to object manipulations. The difference between cell types was not significant in most analyses, possibly due to the low number of active granule cells. It is possible, however, that mossy cells are more responsive to object manipulations than granule cells. An object learning task causes mossy cells, but not granule cells, to increase their phase coupling to gamma oscillations associated with LEC inputs.^50^ Mossy cells, but not granule cells, also show increased c-Fos expression levels following novel object exposure,^51^ and therefore may be more likely to respond to object manipulations, even with familiar objects. Recent studies have reported how tactile cues affect DG activity during learning in head-fixed mice.^30, 31^ While mossy cells encoded cue locations throughout learning, cue-associated activity in granule cells was more common early in training.^31^ The well-trained task used in the present study may therefore reduce the prevalence of object-related granule cell activity.

Alternatively, it may simply be easier to detect object responses in mossy cells. Since we typically observed object-related responses in individual fields, the multiple place fields of mossy cells would increase the odds of detecting an object-related response even if all granule cell and mossy cell firing fields have the same likelihood of responding to a specific object manipulation. Granule cells may also form a sparse representation of nonspatial information, similar to their sparse coding for space.^33, 52, 53^ As mossy cells receive inputs from > 100 granule cells,^35^ mossy cells could provide a convergent readout for the sparse object-related activity of many granule cell inputs.

### Conjunctive encoding of spatial and nonspatial information

The DG is believed to be necessary for the conjunctive encoding of the allocentric context (from MEC) and egocentric content (from LEC) of an experience into a cohesive representation.^14–16, 28, 29, 54, 55^ Behavioral studies have provided support for the role of the DG in forming complex associations between objects, sensory cues and spatial locations.^21, 25, 26, 54, 56^ However, it is unclear how the activity of cells in the DG may contribute to the formation of these conjunctive representations. Conjunctive encoding could be an emergent property of the DG as a whole, arising from distinct populations of dedicated DG “place” cells and “object” cells that process spatial and non-spatial information in parallel,^32^ which is later integrated in downstream regions of the hippocampus.^4, 57^ However, there is already considerable crosstalk between spatial and non-spatial inputs to the hippocampus, within and upstream of the entorhinal cortex.^58, 59^ In the present study, object manipulation affected the activity of different DG cells in each environment; the same DG cell that had a clear object response in one environment could be unaffected by the same manipulation in another environment. This result suggests that the population of object-responsive cells is dynamic and can be reassigned in a context-dependent manner. Single cells could also exhibit changes in activity in response to one object manipulation, but not another, within the same environment. Together, these results indicate that there is not a dedicated population of “object cells.” Instead, our data suggest that DG cells represent both spatial and non-spatial information and can be flexibly deployed to represent different degrees of object and/or spatial information in each environment.

Another possibility for conjunctive encoding is that of a “conjunctive cell” that only fires when multiple conditions are met, such as grid x head direction cells that fire when the animal is in a specific spatial location and facing a specific direction.^60^ By this definition, a DG cell that was similarly responsive to both objects and place might exhibit a unique pattern of firing for each configuration of objects and environments, suggesting that small changes, such as object rearrangement, could induce global remapping across a population of cells with purely conjunctive responses. Instead, we observed more localized changes in the activity of a subset of place fields, within a stable spatial map of the broader environment.

At the level of individual fields, we also typically do not see a purely conjunctive representation. Rather than representing a specific object and the specific spatial location at which it was encountered, most individual fields appear to be associated with the object at multiple spatial locations within the environment and the majority of the object-related responses we observed also track with the object location. However, fields that appear or disappear and are only present in a particular configuration could reflect a purely conjunctive representation. Together our results suggest that conjunctive encoding can occur in single cells through the response of individual fields to object manipulations and multiplexing of object and spatial information within the same cell.

Although individual fields in a subset of DG cells are modulated by object manipulations, we observed global remapping across the population upon exposure to a second environment. Thus, there appears to be a threshold for DG responses wherein small local changes can induce field-specific modulation of spatial firing, but changes in the spatial context induce global remapping, as well as the recruitment of a new set of object-associated cells. Taken together, these data reveal flexibility in the subset of cells that are object-responsive in different contexts, the type(s) of object responses in single cells across sessions, and the response of individual fields to object manipulation within single cells. Our results suggest that spatial and nonspatial information can be integrated and dynamically modulated in response to environmental manipulations by the DG.

Place cells have been proposed to provide a spatial framework for an internal cognitive map.^10^ Information about events and experiences can be encoded onto that spatial map to guide spatial navigation and episodic memory.^10^ To distinguish between experiences in different places, it is essential to encode each distinct environment as a separate spatial map with minimal overlap. However, within the same spatial environment, cues and objects can change over time. Different experiences in the same environment will have slightly different cues, and the ability to flexibly represent such changes without a complete reorganization of the spatial representation is essential. We found the DG population formed stable spatial representations while flexibly responding to object manipulations, which would support the encoding of unique experiences within the same environment.

## Acknowledgments

Supported by grants from the National Institutes of Health (R01NS039456 to J.J.K., R35NS116843 to H.S., R35NS097370 to G-l.M.). We thank Francesco Savelli for feedback on the manuscript.

## Author contributions

D.G., J.J.K., and K.M.C designed the experiments; D.G. and S.H.K collected the data; D.G, K.M.C., S.H.K., and V.P. analyzed the data; H.S, G-l.M., J.J.K, and K.M.C provided supervision and funding; D.G and K.M.C wrote the initial draft of the manuscript; and all authors provided comments and feedback on the manuscript.

## STAR METHODS

### Key resources table

| **REAGENT or RESOURCE** | **SOURCE** | **IDENTIFIER** |
| --- | --- | --- |
| VECTASHIELD Mounting medium with DAPI | Vector Laboratories | Cat#H-1500 RRID AB_236788 |
| Matlab | Mathworks | https://www.mathworks.com/products/matlab |
| Digital Lynx SX data acquisition system | Neuralynx | Neuralynx.com |
| Cheetah acquisition software | Neuralynx | Neuralynx.com |

### Resource availability

*Lead contact:* Requests for further information, data, and other resources can be directed to the lead contact, Kimberly M. Christian

*Materials availability:* This study did not generate new materials

*Data and code availability:* All data and analysis code are available upon request to corresponding authors.

### Experimental Model and Subject Details

We recorded from 21 Tbr2-CreER^T2^::ChR2^f/f^ mice^61^ (13 male, 8 female, 19-29 g, 10-23 weeks old, C57BL/6 background). These mice were used to express Channelrhodopsin (ChR2) in a birth-dated population of adult-born immature granule cells (for a different study). Mice were implanted with a custom-designed recording drive,^62^ consisting of seven independently movable tetrodes, one movable reference tetrode, and one fixed 200 µm diameter optic fiber. The optic fiber was used for another experiment and was targeted to end above the CA1 pyramidal cell layer (1-1.5 mm below the brain surface). Mice were group-housed prior to surgery and individually housed after surgery on a 12 hr light/dark cycle with ad libitum access to water. All surgeries and animal procedures complied with National Institutes of Health guidelines and were approved by the Institutional Animal Care and Use Committees at Johns Hopkins University and the University of Pennsylvania.

### Method Details

#### Surgical procedures

During surgery, mice were anesthetized with isoflurane or a mixture of ketamine/xylazine/acepromazine (70-100, 5-12, and 1-3 mg/kg, respectively). A surgical plane was maintained with supplemental ketamine/xylazine/acepromazine or with isoflurane (to effect). The skull was exposed and cleaned, and a craniotomy was drilled. The dura was cut, and the drive was positioned so that the fiber (center of the diamond-shaped bundle) was located 1.3 mm medial and 2.0 mm posterior to bregma. A ground screw was placed above the cerebellum, and 2-3 additional anchor screws were added to the skull. The craniotomy was sealed with Kwik-Sil (World Precision Instruments), and the drive was secured to the skull with Metabond (Parkell).

#### Training and Behavior

After mice had recovered from surgery, they were food restricted to ∼80-90% of their free-feeding weight and trained to forage for chocolate sprinkles in a series of distinct environments. Mice were trained in 2-4 environments per day (out of 5-6 total trained environments; 1-2 for the present study, 4 for classifier training data and another experiment) for 10-15 minutes per environment. Each environment consisted of a square, octagonal, or circular arena (60-70 cm diameter), with shower curtains surrounding the environment and a prominent polarizing visual cue on the environment walls. The environments were at the same physical location but had different colors, patterns, and textures on the walls and floors, as well as different patterns on the shower curtain surrounding the environment. For 18 mice, a single square environment was used for object recording sessions (63 cm x 63 cm, 30 cm high walls; Figure 1C), and objects were present for all training sessions in this environment. For 3 mice, two square environments were used for object recording sessions (see below). All objects were familiar to the mouse, 4 cm in diameter (at the base), 6 cm tall, and differed in shape, color, and texture. For six mice, a black cylinder and a white rectangular cuboid were used. For all other mice, objects were roughly conical or pyramidal to prevent the top of the object from blocking the recording headstage or tether (Figure 1B).

Once tetrodes reached the DG, object recording days were alternated with days in which animals foraged for food in multiple distinct environments (forage recordings). Data from these forage recording days were used to train a classifier to separate between DG cell types (see below). Forage recording days consisted of five 10-15 minute recording sessions. In the first four sessions, mice foraged for food in an open environment, and each of the four sessions was in a different environment. The final session of the day was a repeated exposure to one of the four environments and was not used for any analysis in the present study.

Object recording sessions consisted of alternating standard (STD) sessions, in which two objects were in a standard configuration, and object-manipulation (MAN) sessions. Manipulation sessions included moved object (MOVE) sessions, in which one of the standard objects was moved to a new location, and added-object (ADD) sessions, in which a third object was added to the environment (Figure 1). Animals foraged for chocolate sprinkles in the environment for 10-15 minutes in each session, and they were removed from the environment between sessions. Each recording day consisted of 2-8 object recording sessions. There was not a fixed number or order of sessions, but different MAN sessions were alternated with STD sessions as long as the mice continued to explore the environments.

In another set of experiments, DG cells were recorded as mice foraged for food in two square environments. One of the two environments was the same square environment used for all other object recording sessions (Figure 1). A second square environment of the same size was also used (Figure 7A). Object locations within this second environment were identical, and the same set of standard objects was used. On each recording day, mice would first forage for food in one of the environments with objects in the STD configuration (STD_A_). Next, one object was moved (MOVE_A_). This pattern of object manipulation was then repeated in the second environment: A STD session (STD_B_) was followed by the same object movement (MOVE_B_). The recording day ended with one more STD session (STD1_A_’ or STD_B_’). All forage and object recording days began and ended with a baseline sleep session, in which mice rested in their home cage for 30+ minutes.

#### Electrophysiological recordings

A total of 366 well-isolated, putative excitatory cells were recorded from the DG of 18 mice as they foraged for food in an environment that contained discrete objects. An additional 112 cells were recorded on tetrodes in the DG of 3 mice as they foraged for food in two square environments. This number of cells may include some cells recorded across days, although object recordings were typically not performed on consecutive days and tetrodes were lowered slowly each day. Tetrodes were made from 12.5 µm nichrome wires (California Fine Wire Co.) and were electroplated with gold to reduce the impedance to ∼200 kOhms. A Digital Lynx SX data acquisition system and Cheetah acquisition software (Neuralynx) were used for recordings. Signals were amplified 1,000–5,000 times and filtered between 600 Hz and 6 kHz (for units) or 1 and 475 Hz (for local field potentials [LFPs]). Spike waveforms above a threshold of 40–80 µV were sampled for 1 ms at 32 kHz, and LFPs were continuously sampled at 1 kHz. Tetrodes were lowered to the dentate gyrus over a period of ∼2-4 weeks. When tetrodes were near the DG, they were lowered < 30 µm per day. In most mice, if cells were present, pulses of blue light were given (following the post-behavior sleep session) to identify adult-born granule cells optogenetically for another experiment. No light pulses were given before behavior on the object recording days analyzed here.

#### Histological procedures

Mice were deeply anesthetized and perfused with phosphate-buffered saline, followed by 4% paraformaldehyde (PFA). The brain was partially exposed and soaked in PFA for 4 hours before tetrodes were retracted and the brain was removed from the skull and drive. The brain was soaked overnight in PFA then transferred to a 30% sucrose solution until the brain had sunk. The brain was then sliced into 40-50 µm thick coronal sections, mounted, and stained for DAPI (Vector Laboratories). Sections were viewed and photographed using a confocal microscope (Zeiss), and tetrode tracks were identified.

#### Unit isolation

Single units were isolated offline using custom-written cluster-cutting software (Winclust, J. Knierim). Multiple waveform characteristics, including spike amplitude, peak, and energy, were used to isolate cells. The isolation quality of each unit was rated on a subjective scale from 1 (very good) to 5 (poor), depending on the separation of the cluster from other clusters and background noise. Isolation was judged independent of behavioral firing correlates, and only cells rated as fair or better (categories 1, 2, and 3) were included in the analysis.

#### Rate maps and place fields

The position of the mouse was monitored by using an overhead camera to track red and green LEDs on the recording headstage. The image was divided into a 64 x 48 pixel rate map where each bin/pixel was a ∼1.3 x 1.3 cm square. The number of spikes when the animal was in each bin of the map was divided by the amount of time the mouse spent in that bin to determine the average firing rate in each bin. This map was then smoothed using an adaptive binning procedure^63^ and spatial information scores were calculated from these rate maps.^63^ A p-value for the spatial information score was determined by a shuffling procedure in which the spike train and location of the mouse were shifted by a random amount (minimum 30 seconds, with the offset points at the end of the data steam wrapped around to the beginning), and the rate map and spatial information score were recalculated. This procedure was repeated 100 times for each cell, and the number of times that the shuffled spatial information score was higher than the observed value determined the p-value for the spatial information score. If the shuffled spatial information score was lower than the observed value for all shuffles, the spatial information score was considered significant (p < 0.01).

Cells were considered active if they had a significant (p < 0.01) spatial information score > 0.5 bits/spike, and had a mean firing rate ≥ 0.1 Hz and (to exclude putative interneurons) < 10 Hz. For active cells, place fields were identified by finding all pixels of the rate map where the firing rate of the cell exceeded 20% of the cell’s peak firing rate. Any group of 30 or more contiguous pixels that passed this 20% threshold was defined as a place field. The center of a place field (place field centroid) was defined as the center of mass of the pixels in that field, weighted by the firing rate of each pixel.

Any cell with a mean firing rate > 10 Hz during the post-behavior sleep session was excluded from analysis (to exclude putative high-rate interneurons). An additional population of cells had a mean firing rate between 2 and 10 Hz, but no significant spatial firing in any environment. These cells may represent a different population of hilar interneurons^33, 64^ and were also excluded from analysis.

### Quantification and Statistical Analysis

#### Cell type classification

Cells were classified as putative granule cells or mossy cells using a random forests classifier.^65, 66^ Random forests is an ensemble learning method in which a large number of decision tree classifiers are generated using a bootstrapped sample of training data and random subsamples of features. Each classifier receives one vote, and the output with the most votes is the final output of the classifier.

The training data used to generate the classifier was collected on separate recording days. On these days, mice foraged for food in four distinct environments (without objects, Figure 2A). A total of 192 putative excitatory cells from 10 mice recorded in forage sessions (without objects) were used to train the classifier for the present study. Only the first four forage sessions from each day were used (the repeated session was excluded). In similar tasks in rats and mice, most granule cells have no place fields in any environment (and were very rarely active in more than one environment) while mossy cells are active in all or most environments.^33, 34^ We therefore could confidently consider cells that did not have place fields in any environment as putative granule cells. Likewise, we considered any cells with place fields in at least three environments as putative mossy cells (Figure 2A). Using this selection criterion, 79 putative granule cells and 62 putative mossy cells were selected. These cells were used as the training data to generate a random forests classifier to separate cell types.

The classifier was trained using firing properties recorded during a post-behavior sleep session. While the spatial firing of cells was used to select putative granule cells or mossy cells for training, no spatial firing features were used for classification. The features used to separate cell types were mean firing rate, burstiness (proportion of interspike intervals that were < 6ms), spike duration (peak to valley of the waveform recorded on the tetrode wire with the largest peak amplitude), and the first principal component of the second derivative of the spike waveform (average waveform of highest amplitude tetrode wire).^34^ The random forests classifier consisted of 300 individual decision trees, and three random features were used at each split. The out-of-bag error rate (an estimate of the classifier’s generalization error) was 5% (Figure S1). This classifier was applied to the post-behavior sleep data recorded on object recording days to identify cells as putative mossy cells and granule cells. While some cells recorded during object sessions may have also been recorded on a previous forage session day, and thus used as part of the training data, tetrodes were moved ∼30 µm daily.

#### Landmark vector analysis

The method used to define landmark vector cells (LVCs) was modified from a previously described method.^4^ For each session, the place field centroids were determined for all place fields in the environment. For sessions with at least two place fields, LVCs were identified using the following procedure. First, vectors connecting each object to each place field centroid were calculated. The difference was then calculated between all pairs of vectors that did not share the same object or the same place field centroid. The pair of such vectors with the smallest difference was determined (see Figure 4A for schematic). Sessions with a minimum vector distance below 7 pixels were considered to be putative LVCs. While a one pixel threshold had been previously used to detect LVCs in the hippocampus,^4^ we used a 7 pixel threshold for two reasons. First, the environment, and thus the size of each pixel, used by Deshmukh and Knierim (2013) was larger: the area of 7 pixels in this study roughly equals the area of 1 pixel from Deshmukh and Knierim (2013). Place fields in mice are also less stable than place fields in rats in certain tasks,^67^ and there are differences in place field stability between DG and other hippocampal subfields.^53^

The distribution of observed minimum vector differences was compared to a random distribution.^4^ To create the random distribution, we first collected all place field centroids from all sessions. Random place field centroids were then assigned to each session. Place field centroids were picked so that the number of randomly assigned place fields equaled the number of recorded place fields in that session, and the distance between all place field centroids was never less than 6 pixels (the minimum distance between place field centroids observed in the data). Vector differences were then calculated for the randomized place field locations (using the same procedure described above) to create the random distribution of minimum vector differences.^4^ This analysis results in a randomized distribution with the same number of sessions (and the same number of objects and place fields in each session) as the real data. To determine if the number of LVCs detected in our data exceeded the number expected by chance, we generated 100 separate random distributions using the procedure described above. For each of these 100 random distributions, the number of sessions with a minimum vector difference ≤ 7 pixels (detected LVCs) was determined. The proportion of random distributions where the number of detected LVCs exceeded the number of LVCs observed in the real data was considered to be the p-value for this comparison. To determine whether our selection of 7 pixels for the LVC detection threshold affected these results, the shuffling procedure was repeated with a 5 pixel threshold (Figure S2A), and with a series of threshold values ranging from 1 to 20 pixels (Figure S2B).

#### Distance to nearest place field

We first identified each MAN session and its preceding STD session and analyzed all pairs of sessions in which at least one place field was present in both sessions. The location of the manipulated object (new object or moved object) in the MAN session was determined (X in Figure 6A, top). The distance from this location to the nearest place field centroid was determined in both the STD and MAN sessions, and the difference between these minimum distances was determined. If a field was near the object location in the MAN session but not in the STD session (e.g., fields appearing at that location or fields moving with the moved object), this difference value would be high, indicating that the place field location was influenced by object manipulation.

As some amount of place field drift and remapping is expected, we repeated the same analysis at the location of the unmanipulated object (Figure 6A). If there was significant remapping or place field drift unrelated to object manipulations, a similar distribution of difference values would be expected at that location. However, if the reduction in distance between the object location and the nearest place field exceeded the 95^th^ percentile of the values obtained using the unmanipulated object location, the reduction was considered to be significant.

#### Rate map correlation analysis

For overall rate map correlation comparisons, all sets of three consecutive sessions that consisted of two STD sessions with a MAN (MOVE or ADD) session between them (STD1 – MAN – STD2) were identified (see Figure 6B for schematic). Only groups of sessions with at least one place field in at least one of the three sessions were analyzed. Rate map correlations were calculated between the rate maps for the STD1 and STD2 sessions (R_STD_) and the STD1 and MAN sessions (R_MAN_). The difference between R_STD_ and R_MAN_ was then calculated and compared to a shuffled distribution (Figure 6B), in which the STD1 map in each group of sessions was replaced with another random STD1 map from another active cell, and the analysis was repeated. This process was repeated 1000 times to generate a shuffled distribution of correlation differences. Cells with correlation difference values that exceeded the 95^th^ percentile of the shuffled distribution were considered to be significant.

“Local rate map” correlations were calculated by first identifying all STD sessions followed by a MOVE session. For each session pair, one object was moved in the MOVE session, and the other object was in the same location in the STD and MOVE sessions. The locations of all objects in the STD and MOVE sessions were determined, and “local rate maps” were created for all object locations in both sessions (STD object locations in STD and MOVE sessions, and MOVE object locations in STD and MOVE sessions). Each local rate map was centered at an object location and consisted of ± 10 pixels in each direction from the object, generating a 21x21 pixel local rate map. The STD rate maps were thresholded to only include activity within place fields (any pixels with a rate < 20% of the peak firing rate were set to 0). The “spatial” rate map correlation was calculated between maps centered at the same spatial location in the STD and MOVE session (the location of the moved object in the MOVE session), and the “object-related” rate map correlation was calculated between maps centered at the location of the moved object in both the STD and MOVE session (Figure 6C). The difference between the “spatial” and “object-related” correlations was calculated. This distribution was compared to the distribution generated by repeating the analysis centered at the location of the unmoved object in each MOVE session. For the “unmoved” object correlations, the “spatial” correlation was calculated as the correlation between maps centered at the location of the unmoved object in both STD and MOVE sessions, and the “object-related” correlation was calculated as the correlation between maps centered at the unmoved object location in the STD session, and the moved object’s new location in the MOVE session (as if the unmoved object were the object that was moved) (Figure 6C). Cells were considered to have a significantly higher “object-related” than “spatial” correlation if the difference between these values exceeded the 95^th^ percentile of the distribution from the “unmoved” object.

#### Subjective response type evaluation

The responses to objects were often complex and did not always conform neatly to the categories described earlier. This complexity made it difficult to derive quantitative measures that captured all of the nuances of the cells’ responses that were apparent to human observers. Thus, to assign putative object responses to individual cells and compare these responses between different environments, we subjectively evaluated object recording sessions. For all active cells, each MOVE session and its preceding STD session were evaluated separately. Only pairs of sessions in which at least one place field was detected in at least one of the two sessions were evaluated. Putative object responses were then assigned to these session pairs.

Three observers (D.G., S.H.K., and K.M.C) separately evaluated the session pairs for object-related responses. Each of the active session pairs was presented in random order. For each active session pair (STD and MOVE), the rate maps, object locations, and firing rates for that pair of sessions was displayed. No other information about the cell or its activity in other recording sessions was presented. The observer would then make a judgment on how object movement had affected the cell’s activity. Cells were classified as 1) no object response/stable spatial firing, 2) Moved and/or rotated field, 3) appear, 4) disappear, 5) trace, or 6) capture (see Figure 5 for examples of each response type). If multiple clear responses could be detected, both responses were recorded. Each observer repeated their evaluation of the session pairs three times (different random order each time), and the response type assigned to the session pair at least 2/3 times was considered their assessment for that session pair. On average, individual observers picked the same response at least two of the three times for 99.6% of session pairs (range 99% - 100%), and the same response was selected all three times for 78% of session pairs (range 75% - 82%).

The subjective response assignments from each observer were then compared. The response assigned by at least two of the three observers was considered the response of that pair of sessions. For 87 of the 95 session pairs (92%), two of the three observers selected the same response type. Session pairs where no two observers selected the same response category were considered ambiguous responses (8 session pairs, 8%).

The probability of observing any specific object-related change in activity was determined across the entire population of active session pairs (in both environments). This probability was used to calculate joint probabilities and estimate how much overlap of object-related activity between environments would be expected by chance.

#### Statistical tests

Statistical tests were calculated in Matlab. Data represent mean ± s.e.m. or median and interquartile range (IQR). The p-values of Wilcoxon rank-sum tests, Wilcoxon signed-rank tests, and χ^2^ tests represent the results of two-tailed tests, and results were considered significant if the p-value was < 0.05. A χ^2^ goodness-of-fit test was used to compare the proportion of cells with object-related activity changes in multiple environments to expected values. The expected values of object-related activity in multiple environments were calculated from the joint probability of detected object-related responses. When comparing observed data to a shuffled or random distribution, cells that passed the 95^th^ percentile of the shuffled or random distribution were considered to be significant at p < 0.05. The proportion of cells that exceeded the 95^th^ percentile was then compared to the null hypothesis that 5% of values would exceed the 95^th^ percentile using a one proportion z-test (test for proportions); p-values for the test for proportions are results of one-tailed tests.

---

**Figure S1.**
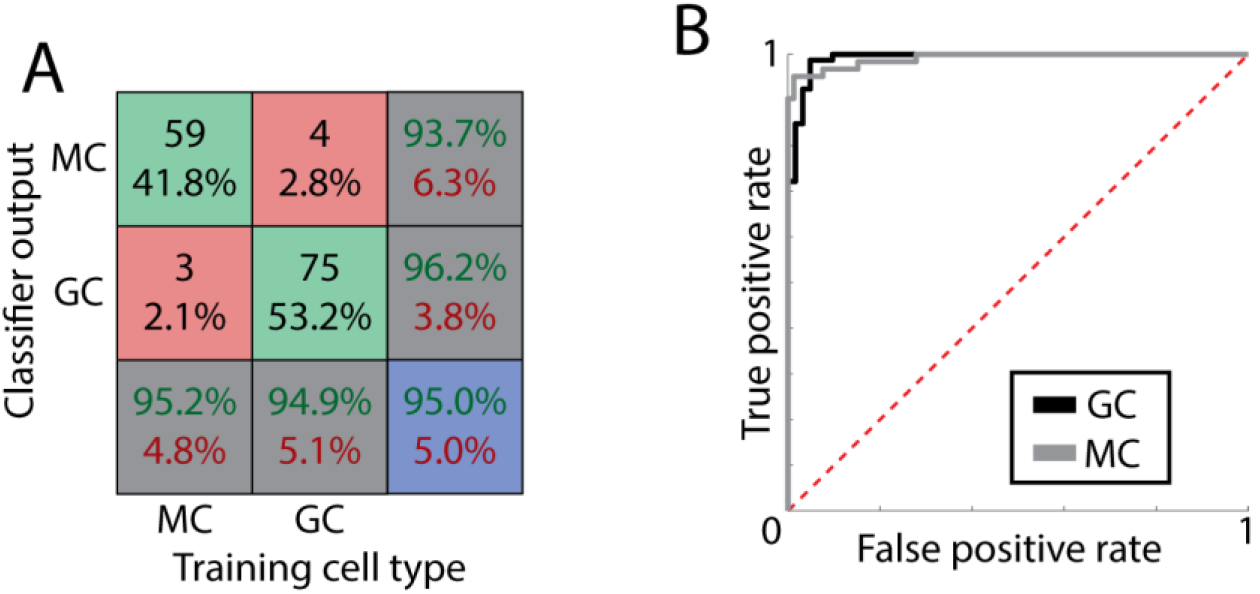
Cell type classification, related to Figure 2. A) Confusion matrix of the out-of-bag error rate (number of cells in training data misclassified by decision trees excluding that cell) for the classifier, comparing cell type label (training cell type, based on the number of active environments) and classifier output. The overall out-of-bag error rate of the classifier was 5%. B) ROC curve for the classification of granule cells (GC, black) and mossy cells (MC, gray). The large areas under the ROC curve for granule cells and mossy cells indicate strong performance of the classifier in discrimination between cell types.

**Figure S2:**
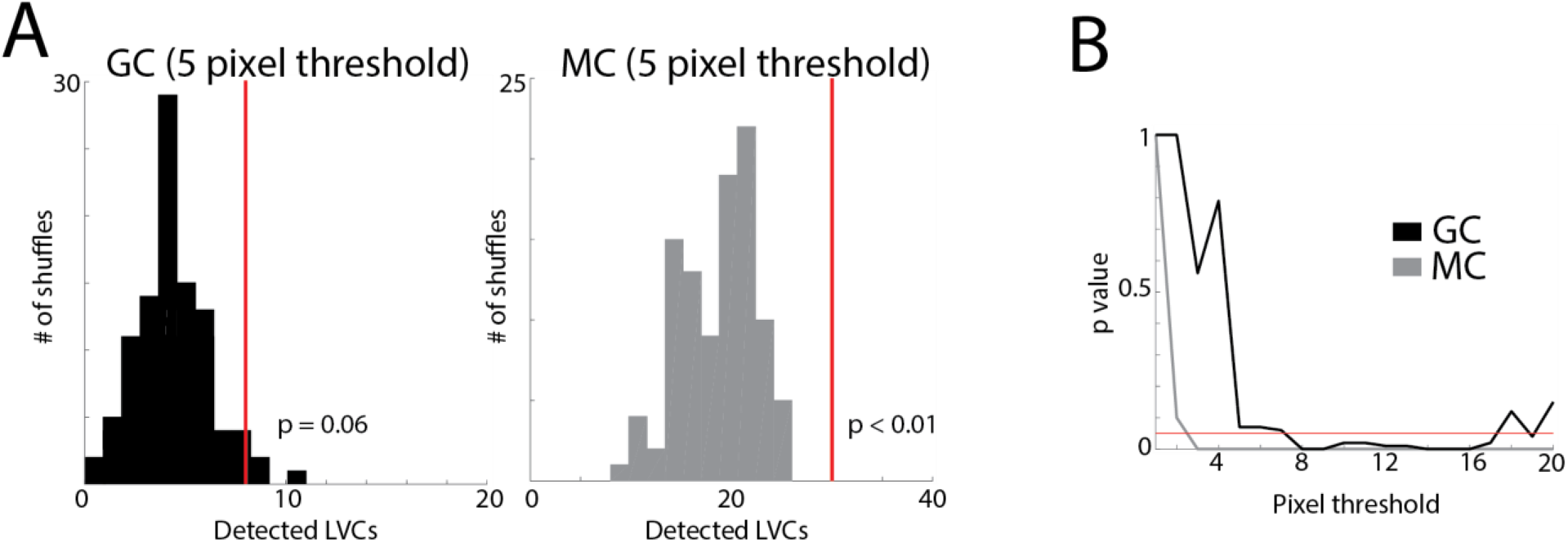
Alternate thresholds for LV detection, related to Figure 4. A) Number of detected LV responses with 5 pixel threshold (maximum pixel difference between vectors to be considered a LV response). Similar to figure 4G, a histogram of the number of detected LV responses in each of 100 random distributions for granule cells (black) and mossy cells (gray) is compared to the actual number of observed LV responses (red line). The number of random distributions with more detected LV responses than observed in the real data is considered to be the p-value. B) The p-value for granule cells (GC, black) and mossy cells (MC, gray) was calculated using pixel thresholds from 1 to 20 pixels. Mossy cells have a significant number of LV responses using any threshold over 2 pixels, while a larger threshold is required for the number of granule cell LV responses to reach significance.

**Figure S3:**
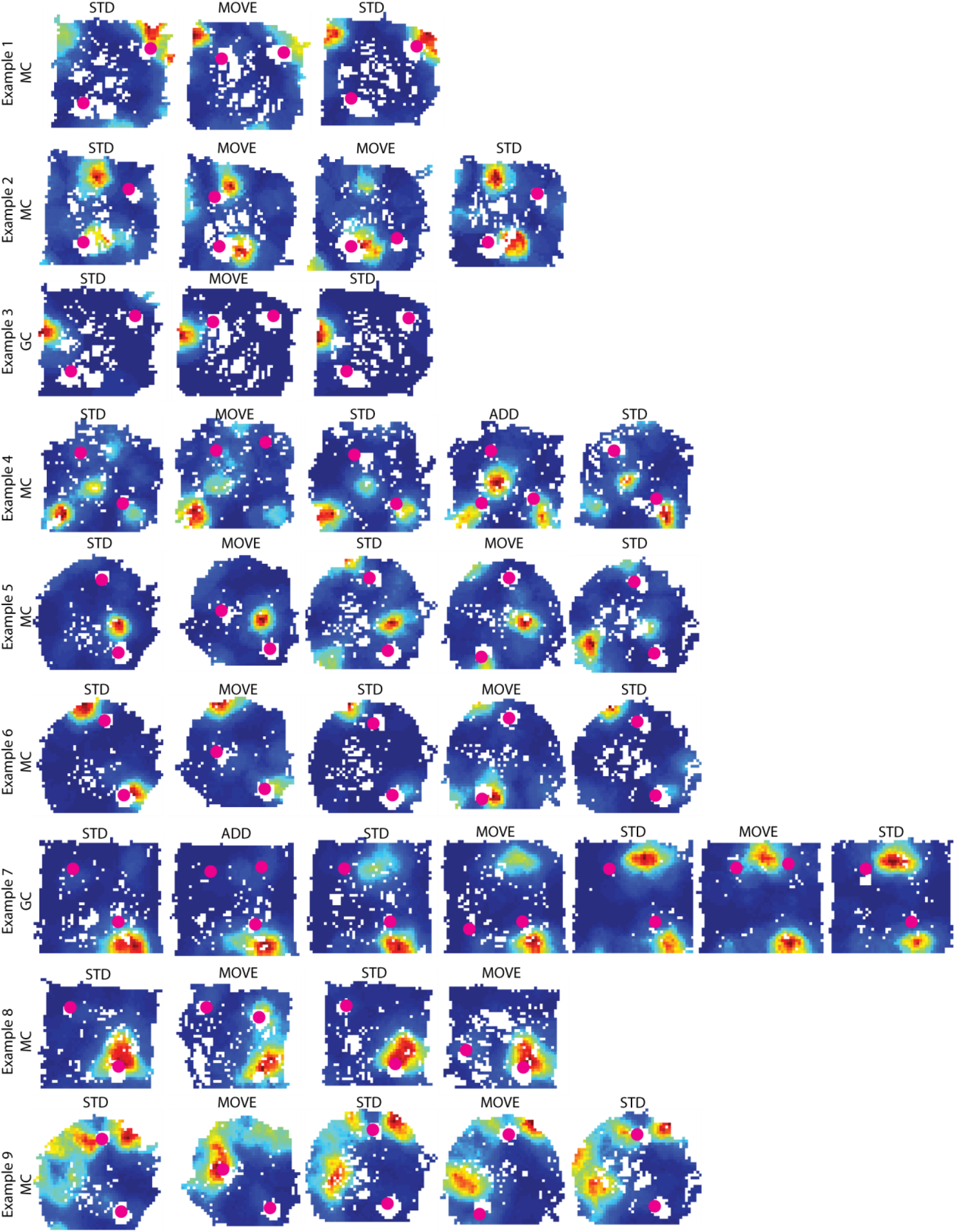
Example DG responses to object manipulation, related to Figure 5. The activity of nine representative cells is shown here, and labeled as a putative mossy cell (MC) or granule cell (GC). Each row is the activity of one cell across all recording sessions in the day. Some cells have a small number of fields that were stable through all sessions and unaffected by object manipulations (examples 1 through 4). This type of activity pattern was the most common; most active DG cells showed stable spatial firing that was unaffected by object manipulations. Example 5 had a single spatial field that persisted through all sessions, while other fields started to emerge in session 3. Example 6 had one field that was unaffected by object movement (top field) and one that moved with the object in session 4 (bottom field). Example 7 had a stable field, as well as a trace response in session 3 that persisted through the remaining sessions. Example 8 shows a cell with a transient response to object movement in session 2 and another stable firing field. Example 9 shows a cell that had more variability in its activity through the sessions, but the field in the top right corner remained relatively stable through all sessions. Together, these cells demonstrate that many DG cells had stable spatial firing, and individual firing fields within single cells could have both stable and object-related activity, highlighting the heterogeneity of DG responses to object manipulation, even within different fields of the same cell.

**Figure S4:**
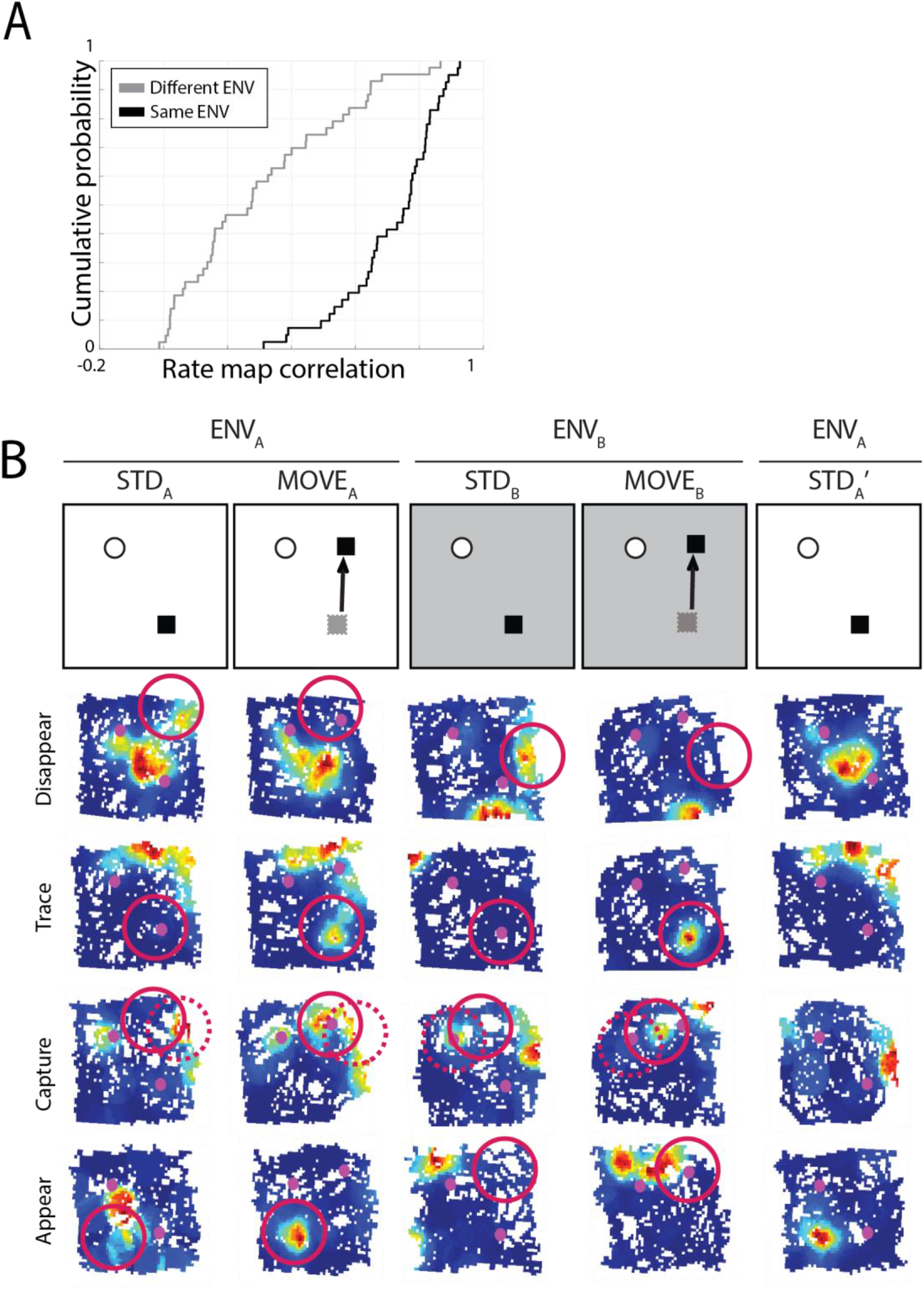
Object-related activity in two distinct environments, related to Figure 7. A) To determine if remapping reported between environments merely reflected rotation of spatial firing patterns, the rate map correlation was calculated between STD sessions in the same or different environments after rotating one map in 90° increments. The maximum value of these rotated rate map correlations was significantly higher for STD sessions in the same environment than different environments (STD_A_ v. STD_B_ 0.30 ± 0.04; STD_A_ v. STD_A_’ 0.71 ± 0.02; rank sum test z = 6.43, p = 1.24 x 10^-10^). B) All cells with the same response to object manipulation in both environments. Each row is the activity of one cell through the five recording sessions. Top to bottom: cell with disappearing field in both environments, field with trace response in both environments, field with capture response in both environments, and cell with appearing field in both environments.

**Table S1:** Statistics for population firing rate comparisons, related to Figure 3. There were no significant differences in the mean firing rate, peak firing rate, number of place fields, or place field size between STD and MAN sessions for granule cells, mossy cells, or all DG cells combined. There was also no difference when only considering ADD sessions (excluding MOVE sessions). STD values for ADD comparisons are slightly different as only cells that had at least one ADD session were included. There was also no difference in the firing rate within 10 pixels of an object (Near) vs. >10 pixels from any object (Away).

|  |  |  |  |  |
| --- | --- | --- | --- | --- |
| **MEAN RATE** | Granule cell: | STD: $0.54 \pm 0.07$ Hz | MAN: $0.51 \pm 0.07$ Hz | signed-rank test<br>$z = 0.67, p = 0.51$ |
| | | STD: $0.60 \pm 0.12$ Hz | ADD: $0.52 \pm 0.09$ Hz | signed-rank test<br>$z = 0.57, p = 0.57$ |
| | Mossy cell: | STD: $0.94 \pm 0.06$ Hz | MAN: $0.88 \pm 0.06$ Hz | signed-rank test<br>$z = 1.62, p = 0.10$ |
| | | STD: $0.90 \pm 0.09$ Hz | ADD: $0.91 \pm 0.10$ Hz | signed-rank test<br>$z = 0.23, p = 0.82$ |
| **PEAK RATE** | Granule cell: | STD: $7.42 \pm 0.97$ Hz | MAN: $7.78 \pm 0.94$ Hz | signed-rank test<br>$z = 0.21, p = 0.83$ |
| | | STD: $7.42 \pm 1.35$ Hz | ADD: $8.88 \pm 1.59$ Hz | signed-rank test<br>$z = 0.51, p = 0.61$ |
| | Mossy cell: | STD: $10.83 \pm 0.66$ Hz | MAN: $10.42 \pm 0.70$ Hz | signed-rank test<br>$z = 1.64, p = 0.10$ |
| | | STD: $10.23 \pm 1.07$ Hz | ADD: $10.19 \pm 1.03$ Hz | signed-rank test<br>$z = 0.23, p = 0.83$ |
| **# OF PLACE FIELDS** | Granule cell: | STD: median 1, IQR 1 – 2 | MAN: median 1, IQR 1 – 2 | rank-sum test<br>$z = 0.12, p = 0.91$ |
| | | STD: median 1, IQR 1 – 1 | ADD: median 1, IQR .5 – 1 | rank-sum test<br>$z = 0.51, p = 0.61$ |
| | Mossy cell: | STD: median 1, IQR 1 – 2 | MAN: median 1, IQR 1 – 2 | rank-sum test<br>$z = 0.64, p = 0.53$ |
| | | STD: median 1, IQR 1 – 2 | ADD: median 1, IQR 1 – 2 | rank-sum test<br>$z = 0.24, p = 0.81$ |
| **PLACE FIELD SIZE** | Granule cell: | STD: $177 \pm 9.7$ pixels | MAN: $176 \pm 13.8$ pixels | rank-sum test<br>$z = 0.60, p = 0.55$ |
| | | STD: $180 \pm 21.7$ pixels | ADD: $190 \pm 33.7$ pixels | rank-sum test<br>$z = 0.09, p = 0.93$ |
| | Mossy cell: | STD: $157 \pm 5.0$ pixels | MAN: $169 \pm 7.2$ pixels | rank-sum test<br>$z = 1.32, p = 0.19$ |
| | | STD: $155 \pm 11.2$ pixels | ADD: $152 \pm 11.7$ pixels | rank-sum test<br>$z = 1.14, p = 0.25$ |
| **NEAR VS. AWAY** | Granule cell: | Near: $0.87 \pm 0.16$ Hz | Away: $0.81 \pm 0.12$ Hz | signed-rank test<br>$z = 0.73, p = 0.46$ |
| | Mossy cell: | Near: $1.33 \pm 0.10$ Hz | Away: $1.28 \pm 0.10$ Hz | signed-rank test<br>$z = 0.98, p = 0.33$ |

**Table S2:** Comparison of observed and expected proportion of cells with object responses, related to Figure 7. We first examined activity in all STD/MOVE session pairs that had at least one place field. We determined the probability of each response type (including session pairs where the spatial firing was unchanged following object manipulation) in all session pairs across both environments (responses detected in ENV_A_ and ENV_B_ were counted separately). Of all active session pairs, there was no detected object response (i.e., stable spatial firing) in 43% of session pairs, and a detected object response in 57% of session pairs. From these probabilities, we calculated the expected proportion of cells that would have the same response in both ENV_A_ and ENV_B._ The expected probability was calculated as the joint probability of the observed proportion of responses in all sessions (left column). If 57% of session pairs have an object response independent of the environment, the probability that the same cell would have an object response in the two tested environments would be 33% (57% of 57%). This expected value was compared to the observed proportion of cells that had the same response in both environments. If a significantly higher proportion of cells had object responses in both environments, this would indicate that a population of DG cells is predisposed to have object-related activity. No significant differences from the expected values were observed, suggesting that there is not a dedicated population of object-responsive cells.

|  | OBSERVED<br>(% OF ALL<br>ACTIVE<br>SESSION<br>PAIRS) | EXPECTED<br>CELLS WITH<br>SPECIFIC<br>RESPONSE IN<br>BOTH ENV | OBSERVED (% OF<br>CELLS) | X<sup>2</sup>(1)<br>VALUE | P-VALUE |
| --- | --- | --- | --- | --- | --- |
| **NO OBJECT**<br>**RESPONSE** | 37/87 (43%) | 18% | 4/32 (12.5%) | 0.67 | 0.41 |
| **ANY OBJECT**<br>**RESPONSE** | 50/87 (57%) | 33% | 10/32 (31.25%) | 0.05 | 0.83 |
| **-MOVE/ROTATE** | 6/87 (7%) | 0.05% | 0/32 (0%) | 0.15 | 0.69 |
| **-APPEAR** | 14/87 (16%) | 2.6% | 1/32 (3.125%) | 0.04 | 0.85 |
| **-DISAPPEAR** | 14/87 (16%) | 2.6% | 1/32 (3.125%) | 0.04 | 0.85 |
| **-TRACE** | 7/87 (8%) | 0.06% | 1/32 (3.125%) | 3.04 | 0.08 |
| **-CAPTURE** | 9/87 (10%) | 1.1% | 1/32 (3.125%) | 1.28 | 0.26 |

